# Spread of pathological human Tau from neurons to oligodendrocytes and loss of high-firing pyramidal neurons in ageing mice

**DOI:** 10.1101/2021.11.24.469849

**Authors:** Tim J. Viney, Barbara Sarkany, A. Tugrul Ozdemir, Katja Hartwich, Judith Schweimer, David Bannerman, Peter Somogyi

**Affiliations:** Department of Pharmacology, University of Oxford, Oxford, OX1 3QT, UK; Department of Experimental Psychology, University of Oxford, Oxford, OX2 6GG, UK

## Abstract

Intracellular aggregation of hyperphosphorylated Tau (pTau) in the brain is associated with cognitive and motor impairments, and ultimately neurodegeneration. We investigate how human pTau affects cells and network activity in the hippocampal formation of THY-Tau22 tauopathy model mice *in vivo*. We find that pTau preferentially accumulates in deep-layer pyramidal neurons, leading to neurodegeneration, and we establish that pTau spreads to oligodendrocytes. During goal-directed virtual navigation in aged transgenic mice, we detect fewer high-firing prosubicular pyramidal cells but the firing population retains its coupling to theta oscillations. Analysis of network oscillations and firing patterns of pyramidal and GABAergic neurons recorded in head-fixed and freely-moving mice suggests preserved neuronal coordination. In spatial memory tests, transgenic mice have reduced short-term familiarity but spatial working and reference memory are surprisingly normal. We hypothesize that unimpaired subcortical network mechanisms maintain cortical neuronal coordination, counteracting the widespread pTau aggregation, loss of high-firing cells, and neurodegeneration.

## Introduction

The majority of tauopathies, defined by intracellular accumulation of hyperphosphorylated Tau proteins (pTau), are ‘sporadic’, and a minority are inherited, caused by mutations of the microtubule-associated protein Tau (*MAPT*) gene.^1^ Humans express six isoforms in the brain, with 3 containing three repeats (3R) each and 3 others containing four repeats (4R) each arising from splice variants.^1^ These isoforms accumulate in different kinds of neurons and glia in different tauopathies, and can have different spatiotemporal distributions. Alzheimer’s disease, amyotrophic lateral sclerosis, Niemann-Pick disease type C and some types of familial frontotemporal dementia and parkinsonism (FTDP) have a mixture of 3R and 4R Tau. Those with only 3R Tau include Pick’s disease and some types of FTDP. The 4R tauopathies include corticobasal degeneration (CBD), progressive supranuclear palsy (PSP), argyrophilic grain disease, and other types of FTDP.^1,2^ The impact on cognitive and motor functions depend on which brain regions and cell types are most vulnerable to developing pTau aggregates and the extent of neurodegeneration. Deficits in spatial memory – our ability to encode and recall environmental contexts – are strongly associated with cortical pTau-related progression from mild cognitive impairment to Alzheimer’s disease.^3–5^ It is generally accepted that the spread of pTau correlates with severity of neurological symptoms, with age being the greatest risk factor.

Animal models provide a wealth of information about the impact of pTau on various aspects of behavior, cognition, network excitability, and plasticity. Tauopathy models can suffer from progressive motor impairments,^6–8^ making it difficult to perform investigations using physiologically-relevant aged animals. We therefore investigated how aggregation of mutant human 4R Tau affected spatial memory and neuronal coordination in awake behaving mice at different ages using the well-characterized THY-Tau22 mouse line. This line gradually develops pTau inclusions from 2-3 months of age,^9^ and has a less severe phenotype than related lines,^8,10^ ideal for investigating cumulative effects of pTau in aged mice. Importantly, heterozygous transgenic (tg) mice and their non-transgenic (ntg) littermates do not develop significant motor impairments during ageing, enabling behavioral testing and awake *in vivo* electrophysiological recordings. Tg mice display prominent neurofibrillary tangles (accumulation of human pTau in somata) as well as neuropil threads (pTau in other processes such as axons). We found that ageing tg mice – which had normal life expectancy – had hippocampal neurodegeneration, showed a reduction in high firing pyramidal neurons, and exhibited reduced short-term familiarity for spatial cues, with pTau spreading from neurons to oligodendrocytes. Despite these changes, neuronal coupling to theta oscillations, and spatial working and reference memory performance showed no detectable differences. We hypothesize that this is due to unimpaired network mechanisms that govern cortical neuronal coordination.

## Results

### Distribution of pTau in ageing mice

To understand how pTau aggregation affects neuronal and network activity, we first mapped pTau in brains of tg mice at different ages. By 2-3 months old (mo), clusters of AT8 immunoreactive (pTau+) neurons were observed in dorsal prosubiculum (ProS), a region between hippocampal dorsal CA1 (CA1d) and dorsal subiculum (SUBd) ^11,12^ (Fig. 1a). The distal part of CA1d (closest to ProS) and SUBd contained very few pTau+ pyramidal neurons at this age. The intensity of pTau in ProS peaked after 8 mo, with large numbers of pTau+ somata and processes (Fig. 1a,b), corresponding to the time when hippocampal-dependent memory deficits have first been observed.^9,13^ Accordingly, a greater number of pTau+ neurons in CA1d were observed (Fig. 2), spreading towards CA2 and a corresponding increase in SUBd. Intensity of pTau decreased in aged mice (>16mo), likely due to cell loss ^9^ and a lack of pTau+ somata in ProS and SUBd of mice >21 mo (Fig. 1a,b). Based on the combination of changes in ProS and CA1, we define young mice as <8 mo, ageing mice as 8-16 mo, and aged mice as >16 mo.

**Figure 1.**
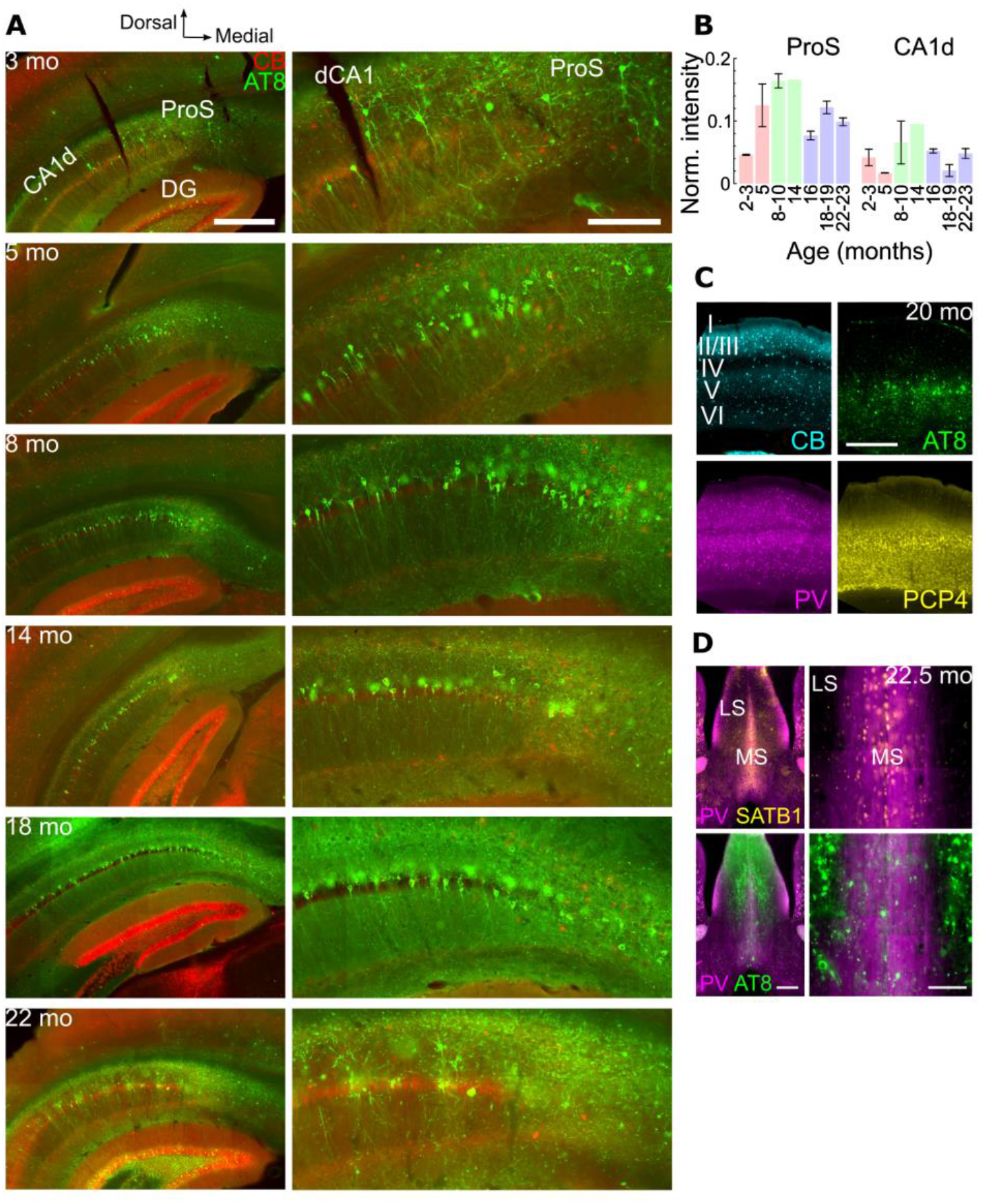
Age-related increase of pTau in THY-Tau22 mice. **(A)** Immunoreactivity for AT8 (pTau+, green) in coronal sections containing the hippocampal formation at different ages (months old, mo). Calbindin (CB, red). Right, enlarged views of distal CA1 (dCA1) and dorsal prosubiculum (ProS). Widefield epifluorescence. **(B)** Normalized intensity measurements of AT8 immunoreactivity (mean, standard error) at 6 different age ranges in ProS and dorsal CA1 (CA1d). Age ranges (left to right): young (pink), 2-3 mo (n=2) and 5 mo (n=2); ageing (green), 8-10 mo (n=2) and 14 mo (n=1); aged (purple), 16 mo (n=2), 18-19 mo (n=2) and 22-23 mo (n=2). **(C)** Immunoreactivity for pTau (green) in deep cortical layers at 20 mo relative to CB (cyan), parvalbumin (PV, magenta) and Purkinje Cell Protein 4 (PCP4, yellow) immunopositive subpopulations. **(D)** Lack of pTau in the medial septum (MS) in a 23 mo tg mouse (case TT21D, female). Processes containing pTau found mostly within lateral septum (LS). Left, epifluorescence tile showing PV (magenta) and SATB1 (yellow) restricted to the MS. Right, enlarged view, note AT8 (green) mostly in LS. Scale bars (μm): A (left), 500; A (right), 200; C, D (left), 500; D (right), 100. Cases in A, top to bottom (mouse, sex): TT114A m; TV132 m; TV143 m; TV136 f; TV139 m; TT33G f.

**Figure 2.**
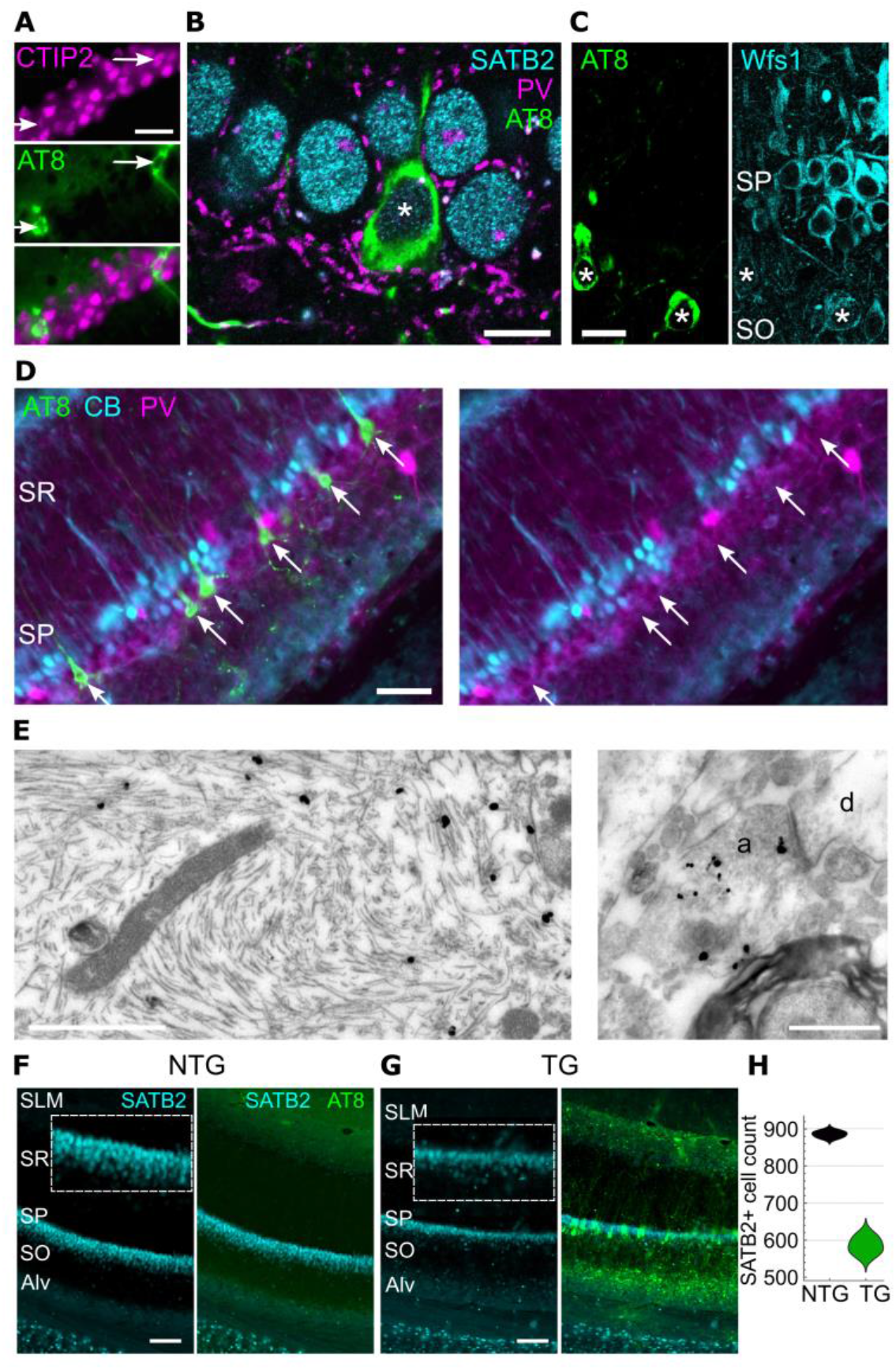
Preferential accumulation of pTau in deep-layer CA1 pyramidal cells. **(A)** Neurons immunoreactive for pTau (AT8, green, arrows) in an ageing mouse show nuclear immunoreactivity for CTIP2 (magenta) in CA1d SP. Widefield epifluorescence, case TV136 (14 mo). **(B)** A pTau+ neuron (asterisk) in distal CA1d SP immunopositive for SATB2 (cyan) in an aged mouse. Note PV+ basket terminals around somata (magenta). Confocal single optical section (0.43 μm thick), case TT21D (23 mo). **(C)** Weak Wfs1 immunoreactivity (cyan) in pTau+ distal CA1d neurons (AT8, green, asterisks) in a young mouse. Confocal single optical section (1 μm thick), case TV133 (6 mo). **(D)** Cells in CA1d SP immunopositive for pTau (AT8, green). Lack of detectable CB (cyan) or PV (magenta) immunoreactivity in pTau+ neurons within deep SP (arrows). Widefield epifluorescence. **(E)** Electron micrographs of AT8-positive filaments (gold-silver particles) within a cell body (left) and an axon terminal (a, right). Dendrite, d. From an aged mouse (case TT33G, 21 mo female). **(F, G)** Immunoreactivity of SATB2 (cyan) and AT8 (green) in CA1d of aged ntg and tg mice (22 mo). Insets, enlarged views of SP. Widefield epifluorescence. **(H)** Distribution chart of SATB2+ nuclei counts in CA1d (n=3 mice/genotype). Alveus, Alv; stratum oriens, SO; stratum radiatum, SR; stratum lacunosum moleculare, SLM. Scale bars (μm): A, C, 20; B, 10; D, 50; E, 0.5; F, G, 100.

In young mice, pTau+ dentate granule cells were rare at the septal pole (sparse pTau: n=4/5 mice; moderate pTau: n=1/5 mice) but common at the temporal pole (moderate or extensive: n=3/6 mice; absent or sparse: n=3/6 mice; Figs. 1a, S1a-c). Mossy fibers showed variable levels of pTau, likely originating from granule cells of the temporal pole (Fig. S1a,c). In ageing mice, the septal pole contained mostly moderate pTau (n=2/3 mice; sparse: n=1/3 mice) and the temporal pole more extensive pTau (n=2/3 mice; moderate: n=1/3 mice). In aged mice, the septal pole was variable (moderate or extensive: n=5/10 mice; sparse: n=5/10 mice) whereas the temporal pole showed extensive pTau (n=6/7 mice; sparse: n=1/6 mice; Figs. 1a, S1a-c). Temporal (ventral) CA1 and CA3 also showed gradual increases in pTau levels during ageing (Fig. S1c).

The most prominent and earliest site outside the hippocampal formation was the basolateral amygdala, exhibiting moderate pTau at 2.5 mo and extensive pTau in all animals >4.5 mo (n=14 mice; Figs. S1a, S2a-c). The medial prefrontal cortex showed modest pTau (Fig. S2a). The entorhinal cortex had similar modest levels of pTau at all ages (Fig. S1d). In isocortical areas, pTau was distributed primarily around layer 5, remaining sparse until much later in life (sparse in n=10/10 young and ageing mice; moderate or extensive in n=7/9 aged mice; Figs. 1c, S1a-c, S2b). In ntg littermates, pTau was not observed (n=16/16 mice, 2-24 mo). Lipofuscin fluorescence was detected in all aged mice. In summary, the distribution in the hippocampal formation, isocortex and amygdala largely confirmed previous reports.^9,13,14^

In contrast to the neuronal pTau distribution described above, key subcortical areas linked to the generation of theta frequency rhythmic activity were unaffected, including anterior and midline thalamus, medial septum (Fig. 1d, S2b,c), mammillary bodies, and supramammillary nucleus. Other areas of the hypothalamus, and basal ganglia, were also largely unaffected (Fig. S1a). In the midbrain, neuronal pTau was found in the red nucleus, deep layers of the superior colliculus, periaqueductal gray, and pontine nuclei, but not in the raphe nuclei, ventral tegmental area, substantia nigra, or interpeduncular nucleus (from n=4/4 aged mice). Midbrain dopaminergic (tyrosine hydroxylase immunoreactive) neurons lacked pTau. Furthermore, pTau+ axons were prominent in fimbria, fornix, lateral septum and thalamic reticular nucleus, most likely originating from affected hippocampal/cortical neurons (Figs. 1d, S1a, S2b,c, 3a,b).

### Deep-layer CA1 pyramidal neurons preferentially accumulate pTau

Hippocampal CA1 is critical for spatial/episodic memory and contains place cells. Neurons containing pTau were immunoreactive for CTIP2, SATB2, and Wfs1, which are markers of CA1 pyramidal cells ^15,16^ (n=50/51 pTau+ SATB2+ CA1d cells, n=3 mice; Fig. 2a-c). Immunoreactivity for Wolframin (Wfs1) was extremely low compared to nearby unaffected neurons and was localized to the plasma membrane (n=11/23 CA1d cells weakly immunopositive, n=12/23 undetectable; Fig. 2c, S3a), suggesting that pTau affects Wfs1 levels.^17,18^

The upper, compact sublayer of stratum pyramidale (SP) primarily consists of calbindin immunopositive (CB+) neurons, and the lower sublayer neurons are mostly CB immunonegative (CB−, lacking detectable levels of CB).^16,19^ Interestingly, the majority of pTau+ CA1 SP neurons were found in the deep sublayer and were CB− (95 ± 6 % for n=3 young mice < 8 mo, n=3/52 CB+ pTau+ cells; 85 ± 5 % for n=3 ageing mice 8-14 mo, n=9/60 CB+ pTau+ cells; 79 ± 11 % for n=5 aged mice 16-23 mo, n=11/59 CB+ pTau+ cells; Fig. 2d). Cells that were pTau+ lacked parvalbumin (PV) and calretinin (CR) immunoreactivity (n=0/36 PV+ or CR+ cells from 2 mice), which mark subpopulations of GABAergic neurons. In the ProS, most pTau+ cells lacked CB (n=0/30 CB+ in n=3 young mice, n=1/30 CB+ in n=3 ageing mice, n=3/47 CB+ in n=5 aged mice). We confirmed pTau in pyramidal neurons by electron microscopy and also localized pTau to axon terminals (Fig. 2e). Thus, pTau preferentially accumulated in CB− pyramidal neurons.

There is previous evidence of a general loss of CA1 cells in tg mice over 12 mo.^9^ To determine whether this was due to a loss of pyramidal neurons, we counted SATB2+ nuclei along CA1d SP; SATB2 was not detectable in CA2 or ProS/SUBd pyramidal neurons. In aged mice, CA1 SP was markedly thinner, particularly towards ProS, where the most abundant pTau inclusions were observed (Figs. 2f-h, 1a,b). There were fewer pyramidal cells in aged tg mice compared to aged-matched ntg mice (34% reduction; mean ± s.d. 886 ± 9 nuclei for n=3 ntg mice versus 585 ± 24 nuclei for n=3 tg mice; *t*_(4)_=20.6, *P*< 0.0001; 22-23 mo; Fig. 2h). In contrast, there was no difference in CA1 SATB2+ nuclei counts in young mice (426 ± 13 nuclei for n=2 ntg mice versus 483 ± 51 nuclei for n=2 tg mice; *t*_(2)_=1.5, *P*=0.2643; 4-6 mo). Interestingly, we did not find a difference in CA1 DAPI+ nuclei counts in ageing mice (659 ± 72 nuclei for n=2 ntg mice versus 555 ± 31 nuclei for n=2 tg mice; *t*_(2)_=1.9, *P*=0.2020; 15-16 mo). Thus, we suggest the accumulation of pTau in pyramidal neurons eventually leads to their degeneration in aged mice.

### Spread of pTau from pyramidal neurons to oligodendrocytes

In aged (n=11/11, 18-27 mo) and ageing (n=5/6, 8-16 mo) tg mice but not in young tg mice (n=7/7, 2-6 mo), we observed coiled bodies – intensely AT8 immunoreactive ‘flame-like’ rings of pTau – in many brain regions, especially in white matter tracts containing pTau+ axons (Figs. 3, S4), such as the fimbria, fornix, alveus, dorsal hippocampal commissure, corpus callosum, and stria terminalis. Coiled bodies were also in hippocampal stratum oriens, subicular complex, granular retrosplenial cortex (Fig. S4), isocortex, dorsal striatum, lateral septum, and were similar in appearance to those found in Alzheimer’s disease.^20^ The anterior thalamus and mammillary bodies had scattered coiled bodies. In the brainstem, the substantia nigra pars reticulata, ventral tegmental area, pontine nuclei and the interpeduncular nucleus contained coiled bodies, a pathology found in 4R tauopathies such as CBD and PSP.^21,22^

**Figure 3.**
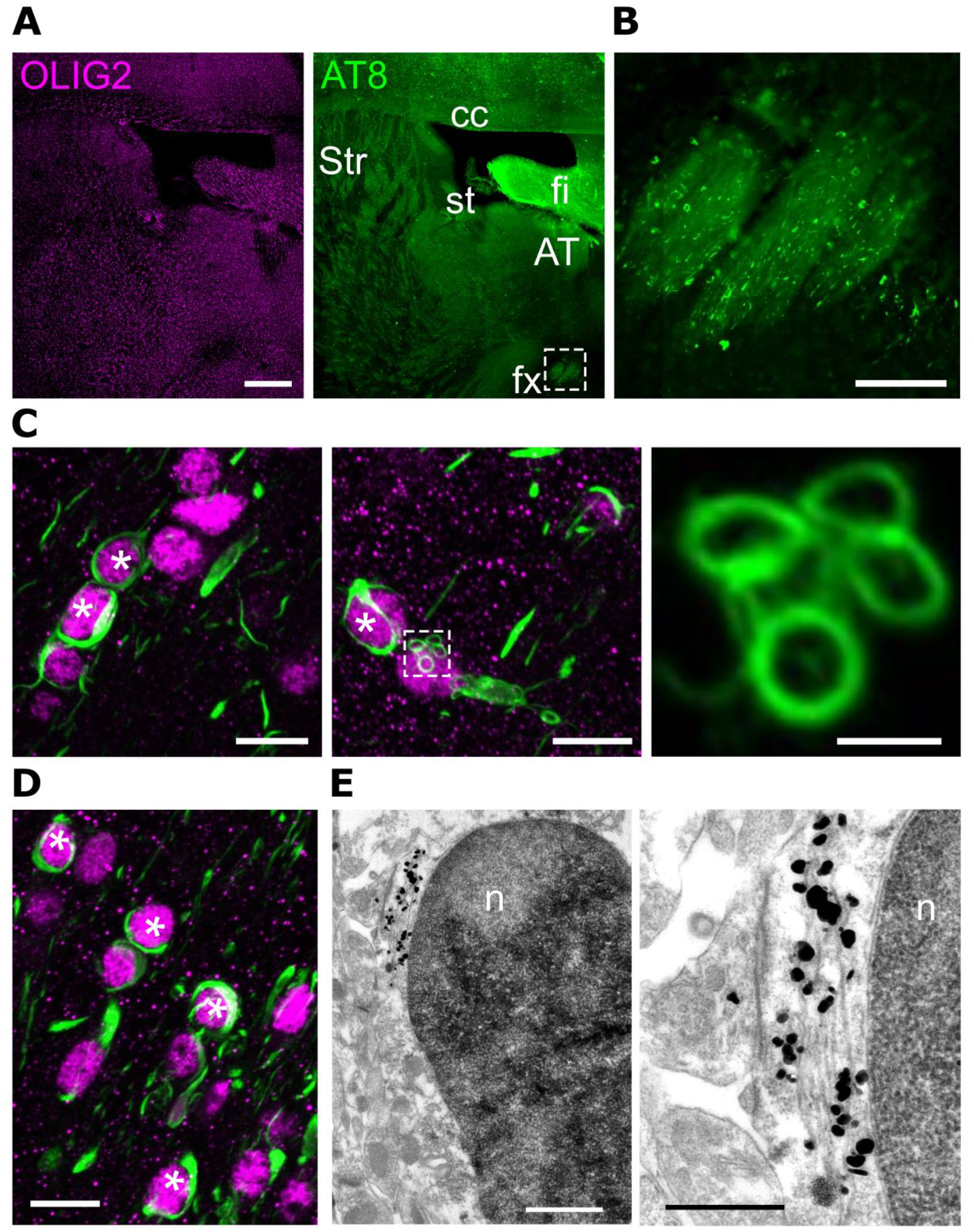
Accumulation of pTau in oligodendrocytes. **(A)** Olig2 (magenta) and pTau (AT8, green) from an aged (20 mo) tg mouse (widefield epifluorescence). Note high density of pTau in fimbria (fi). **(B)** Enlarged view of boxed region from A of the fornix (fx), showing neuropil threads (axons) and coiled bodies. **(C)** Nuclear immunoreactivity for Olig2 (magenta, asterisks) surrounded by coiled bodies (green, AT8). Right, enlarged view of ‘microcoils’ near an Olig2+ nucleus (boxed region, middle panel). **(D)** Coiled bodies colocalizing with Olig2+ nuclei in the fimbria. **(E)** AT8-positive filaments (gold-silver particles) in the cytoplasm of an oligodendrocyte in the ProS next to an Olig2-immunolabeled nucleus (n) (peroxidase labeling) from an aged mouse (case TT21D, 23 mo). Right, higher magnification. Abbreviations: Str, dorsal striatum; cc, corpus callosum; st, stria terminalis; AT, anterior thalamus. Scale bars (μm): A, 500; B, 100; C, 10, 10, 2; D, 10; E, 0.5. Confocal maximum intensity z-projection thickness (μm): C, 4, 9.6, 2.4; D, 4.

Coiled bodies colocalized with Olig2, an oligodendrocyte transcription factor (Figs. 3, S4e,f). Smaller coils (< 2 μm diameter) associated with Olig2-immunoreactive nuclei (Fig. 3c) may represent densely packed pTau. Olig2 was not observed in pTau+ neurons (Fig. S4e). We confirmed the presence of pTau+ filament bundles in oligodendrocytes using electron microscopy (Fig. 3e).^23,24^ To our knowledge, the Thy1 promoter is not active in oligodendrocytes.^25^ As coiled bodies were not observed in younger mice, despite accumulation of pTau in neurons at all ages investigated (Fig. 1), we suggest pTau spreads from neurons to oligodendrocytes via pTau-containing axons (Fig. 3).

### Reduced short-term familiarity but unimpaired spatial reference and working memory

To test behavioral effects of pTau accumulation and neurodegeneration, we assessed short-term hippocampal-dependent spatial memory (age range, 13-24 mo; median [interquartile range] 15 [4] mo ntg, and 14 [5] mo tg mice) using a spontaneous spatial novelty preference Y-maze task (Fig. 4a, STAR Methods). The total distance mice covered in the exposure phase was similar for both genotypes (mean 1.78 ± s.d. 0.61 m for n=18 ntg mice versus 1.75 ± 0.45 m for n=15 tg mice; *t*_(31)_=0.1635, *P*=0.8712; Fig. 4a), indicating tg mice were not hyper- or hypoactive. Discrimination ratios (time in novel over total time in novel and other arms) were significantly different (0.64 ± 0.10 for n=22 ntg mice versus 0.52 ± 0.16 for n=20 tg mice; *t*_(40)_=2.9637, *P*=0.0058; Fig. 4a). This was explained by ntg mice spending more time in the novel arm compared to chance (*t*_(21)_=6.6435, *P*< 0.0001; 110 ± 27 s novel arm, 61 ± 22 s other arm), as expected for wild type animals.^26^ Tg mice failed to discriminate (*t*_(19)_=0.4840, *P*=0.6339; 84 ± 25 s novel arm, 85 ± 45 s other arm), consistent with previous studies reporting spatial novelty preference impairment in younger tg mice.^27,28^

**Figure 4.**
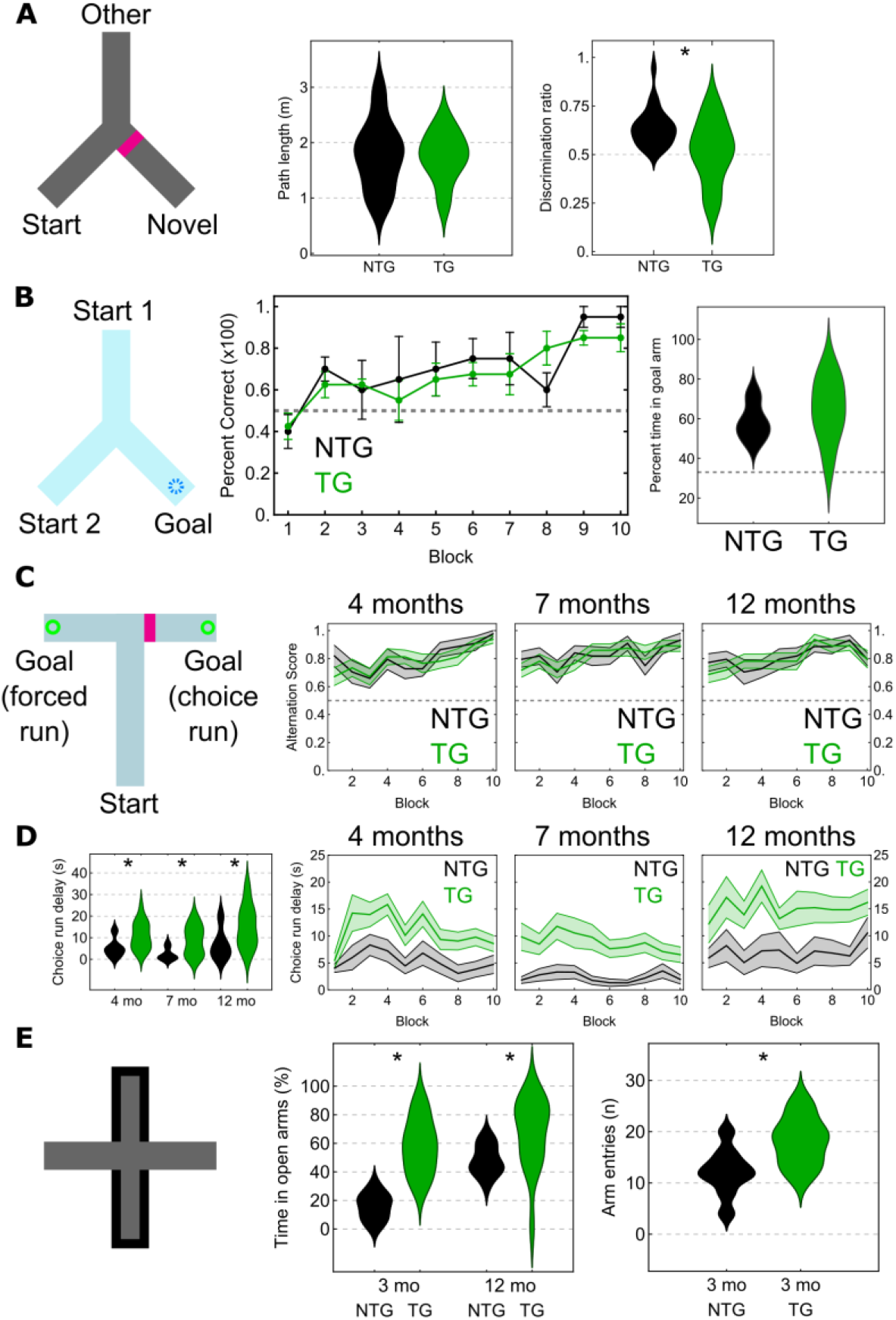
Selective impairments in spatial novelty preference. **(A)** Short-term spatial novelty preference Y-maze test. Left, schematic (extra-maze cues not shown). Novel arm is blocked off for exposure phase and block removed for test phase (magenta bar). Middle, total distance traveled during test phase. Right, distribution chart for discrimination ratio; dashed line, no discrimination (0.5, chance). **(B)** Water escape Y-maze task. Left, schematic: two possible start arms and a goal arm containing a hidden platform submerged in opaque water. Middle, mean values for % correct over time (blocks). Error bars, standard error. Right, distribution chart for probe test. Dashed lines, chance. **(C)** Non-matching to place T-maze task. Left, schematic: start arm and rewarded goal arms. In choice run, block (magenta bar) is removed. Mean alteration scores over time (blocks) for each age. Error bands, standard error. Dashed line, chance. **(D)** Left, distribution chart of mean choice run delay times (ntg, black; tg, green) during training in T-maze at 4, 7, and 12 mo. Right, mean and standard error across blocks. See Fig. S5b for forced run delay times. **(E)** Elevated plus-maze with open and closed arms (left). Distribution charts for % time in open arms (middle) and number of arm entries (right). Asterisks in A, D, E: significant differences. Abbreviations: mo, months old; NTG, non-transgenic; TG, transgenic mice.

To evaluate spatial reference memory, we tested 14 mo mice in a hippocampal-dependent water escape Y-maze task (Fig. 4b, STAR Methods).^29–31^ Mice of both genotypes learned the location of the hidden platform. Tg mice performed similarly to ntg littermates during acquisition training (choice accuracy, repeated-measures two-way ANOVA: genotype *F*_(1,9)_=0.1123, *P*=0.7452; training block *F*_(9,81)_=6.809, *P*=0.0026; genotype x training block, *F*_(9,81)_=0.9458, *P*=0.4908; Fig. 4b; one ntg mouse was excluded as it failed to make choices after day 3). On day 11, we removed the hidden platform and let mice swim freely for 60 s. We found no difference (58.6 ± 9.68% of time in the goal arm for n=4 ntg mice vs 64.56 ± 15.96%, n=7 tg mice; *t*_(9)_=0.6660, *P*=0.5221; chance is 33.3% in the goal arm; Fig. 4b). In summary, based on this task we could not detect deficits in spatial reference memory.

We tested spatial working memory in a non-matching to place T-maze task (Fig. 4c, STAR Methods).^32^ We tested mice at 4, 7 and 12 mo (n=11 ntg and n=16 tg mice; within-subject, longitudinal design) under mild food restriction to 90.7 ± 2.7% of their free-feeding weight. Behavioral performance improved with training (alternation score, repeated-measures ANOVA: training block *F*_(9,225)_=8.497, *P*<0.001; Fig. 4c), independent of age and genotype (age *F*_(2,50)_=0.534, *P*=0.590; genotype *F*_(1,25)_=0.295, *P*=0.592; age x genotype *F*_(2,50)_=0.024, *P*=0.976), with an overall similar performance across blocks (age x block x genotype *F*_(18,450)_=0.522, *P*=0.948; Fig. 4c). Improvements in correctly alternating on the choice run occurred at all ages (Fig. 4c). All training was carried out within the same context; a new context would require mice to form new spatial maps, which might reveal differences in genotypes. One cohort (n=6 ntg and n=9 tg mice) was re-tested at 13 mo in a different context (STAR Methods). Mice approached a high alternation score by the 5^th^ block for both genotypes (Fig. S5a).

During T-maze training we noticed that many mice would wait at the end of the start arm before voluntarily initiating a run. Strikingly, tg mice waited significantly longer than ntg mice before initiating their choice run (4 mo: 4.6 [3.9] s for ntg mice, 10.2 [11.1] s for tg mice, *U*=41, *P*=0.0218; 7 mo, 1.0 (4.1) ntg, 9.3 (12.3) s tg, *U*=23.5, *P*=0.0016; 12 mo, 5.1 (7.9) s ntg, 13.5 (14.0) tg, *U*=38, *P*=0.0145; Fig. 4d). However, this was also the case for the forced run (4 mo, 1.8 [1.4] s ntg, 3.8 [3.5] s tg, *U*=40, *P*=0.0191; 7 mo, 0.5 [1.3] s ntg, 3.2 [5.3] s tg, *U*=29.5, *P*=0.0042; 12 mo, 1.5 [3.1] s ntg, 6.9 [12.0] s tg, *U*=34.5, *P*=0.0089; Fig. S5b), suggesting the delay in choosing is not directly related to making decisions. Although ntg and tg mice had similar delay times in the first session (block 1), they rapidly diverged over subsequent days (Figs. 4d, S5b). Once mice left the end of the start arm, actual run times were similar, suggesting unimpaired ambulation (Fig. S5c). In summary, although we did not detect a spatial working memory impairment in this task, tg mice took longer to initiate both forced and choice runs.

Mice were also placed in an elevated plus maze, an ethological test of unconditioned anxiety. Wild type rodents tend to spend more time in closed than open arms, and time in open arms is increased in rats with ventral hippocampal lesions;^33,34^ this region was also enriched with pTau (Fig. S1c). In young (3 mo) mice, tg mice spent a significantly greater proportion of time in open arms than ntg mice (17.1 [13.7] % of time in open arms for n=9 ntg mice versus 58.3 [29.9] % for n=5 tg mice; *U*=45, *P*=0.0022), consistent with previous reports at 6 mo.^9^ There was also a slight difference in arm entries (12 [4.25] entries for 9 ntg mice versus 19 [7] for 5 tg mice; *U*=8, *P*=0.0448; Fig. 4e). In a separate cohort of 12 mo mice that had been extensively handled and tested in other behavioral tasks but were naïve to the plus maze, ntg mice (n=11) spent 46.9 [16.4] % of time in open arms compared to 75.2 [34.0] % for tg mice (n=16) (*U*=142, *P*=0.0083; Fig. 4e). Taken together, we conclude that pTau aggregation affects short-term familiarity and anxiety-related activity but leaves spatial reference and working memory unimpaired.

### Network oscillations are preserved in the presence of hippocampal pTau

To determine whether intracellular pTau affects neuronal and network activity, we used glass electrodes to record network oscillations and single neurons in the hippocampal formation of both awake head-fixed and freely moving mice. We first focused on 5-12 Hz theta oscillations, which reflect coordination of neuronal activity ^35–38^ required for spatial memory. Prominent theta oscillations were observed in aged (17-24 mo) head-fixed mice while they ran on a running disc (Fig. 5a), with similar peak theta frequency values across genotype for slow speeds (4-6 cm/s; 6.35 ± 0.59 Hz for n=3 ntg mice; 6.70 ± 0.45 Hz for n=3 tg mice; *t*_(4)_ =0.947, *P*=0.3975) and faster speeds (6-10 cm/s; 6.70 ± 0.45 Hz ntg; 7.76 ± 0.60 Hz tg; *t*_(4)_=2.454, *P*=0.0701). In a different context involving trained water-restricted aged mice (22-24 mo) running on an air-suspended jet ball along a virtual linear track, the theta peak was also similar for both genotypes (6.80 ± 0.16 Hz for n=3 ntg mice; 6.38 ± 0.27 Hz for n=3 tg mice; *t*_(4)_=2.324, *P*=0.0808; Fig. 5a). These data suggest that the sampled theta frequency oscillations are unaffected by pTau.

**Figure 5.**
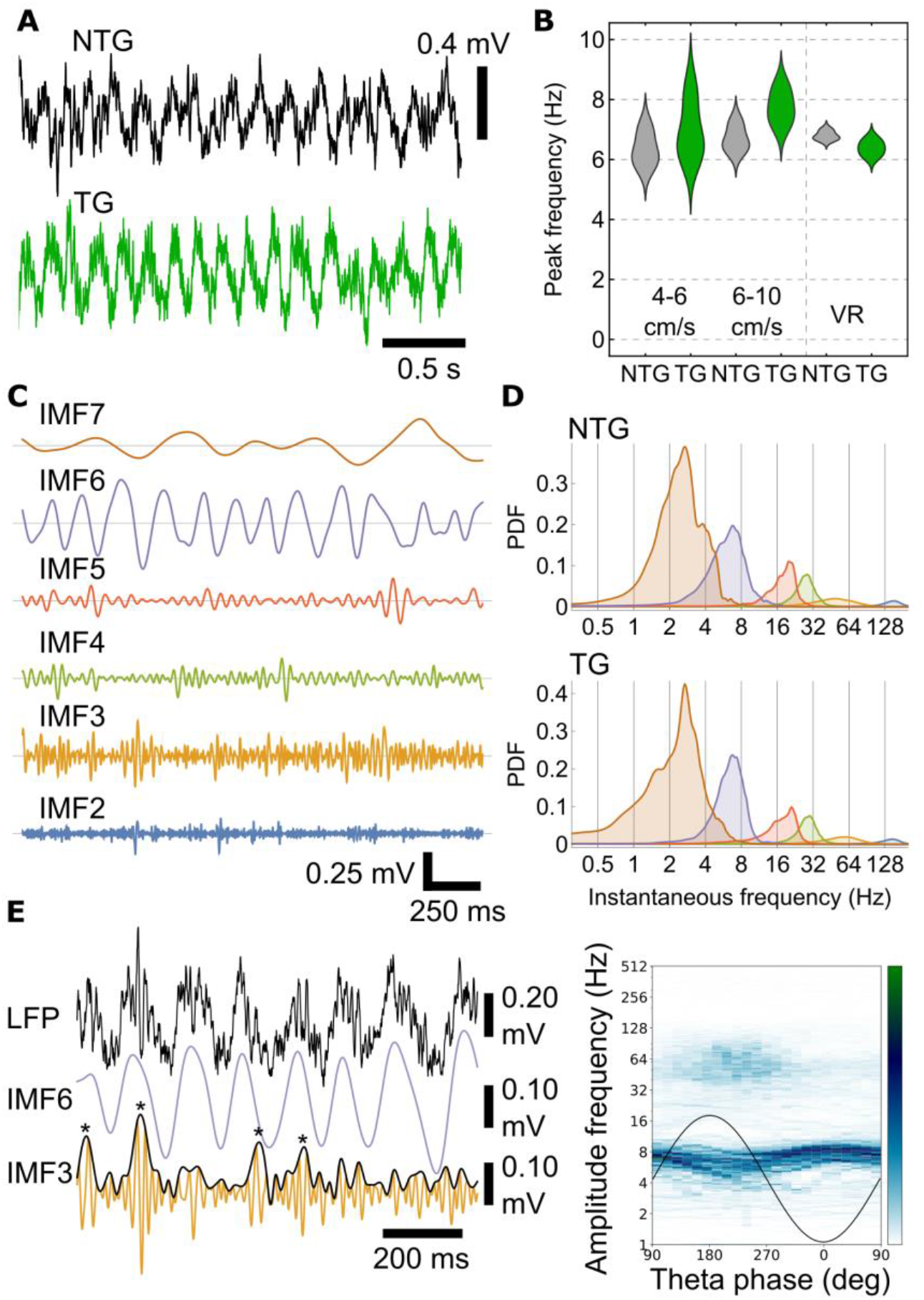
Similar hippocampal network oscillations in tg and ntg mice. **(A)** LFP during movement for aged littermates (ntg mouse, black, case TV140; tg mouse, green, case TV139; 18 mo). **(B)** Distribution chart of peak theta frequencies for two speed bins during spontaneous movement on a running disc, and during navigation in virtual reality (VR), for aged mice. **(C)** Empirical mode decomposition of glass electrode LFP in CA1d SP during movement (aged ntg mouse, 23 mo, case TV147). Theta (IMF6), slow-gamma (IMF4), mid-gamma (IMF3) and fast-gamma (IMF2) oscillations; IMF7 likely corresponds to ‘delta’ and IMF5 to ‘beta’. **(D)** Probability density functions (PDF) of instantaneous frequency for the 7 IMFs. Top, aged ntg mouse (23 mo, case TV147). Bottom, aged tg mouse (18 mo, case TV139). Note similar distributions and spectral peaks. **(E)** Left, unfiltered wideband LFP from an ageing tg mouse (13 mo, case TV136) with IMF6 (purple, theta) and IMF3 (orange, mid-gamma). Black curve, instantaneous amplitude for IMF3; note higher amplitude near peaks of some theta cycles (asterisks). Right, co-modulogram of a 30 s movement epoch binned for IMF6 (theta) phase. Note phase amplitude coupling for mid-gamma (IMF3) around peak-to-descending theta phase. Theta frequency also modulated by phase. Black curve, sinusoidal schematic theta cycle. Color bar: green, maximum; white, minimum amplitude.

In order to investigate potential network changes more closely, we compared the spectral content of local field potentials (LFPs) using empirical mode decomposition (EMD), which is more appropriate for non-linear non-stationary signals.^39,40^ In aged mice (18-24 mo), we sampled LFPs recorded with glass electrodes and silicon probes for movement periods as above and used masked EMD to obtain intrinsic mode functions (IMFs; STAR Methods). Theta corresponded to IMF6 (Fig. 5c) with a mean instantaneous frequency of 7.3 ± 0.4 Hz in ntg mice (n=6) and 6.9 ± 0.2 Hz in tg mice (n=5). We detected slow-gamma (IMF4), mid-gamma (IMF3) and fast-gamma (IMF2) oscillations (Fig. 5c,d), with similar distributions for both genotypes (IMF4, 28.7 ± 0.7 Hz for n=6 ntg mice, 29.3 ± 1.2 Hz for n=5 tg mice; IMF3, 56.8 ± 0.8 Hz for n=5 ntg mice, 59.3 ± 4.1 Hz for n=4 tg mice; IMF2, 140.1 ± 6.3 Hz for n=6 ntg mice, 146.4 ± 4.3 Hz for n=5 tg mice; Figs. 5d, S6a).

Higher frequency oscillations are typically coupled to phases of slower oscillations in rodents and humans.^39,41^ Impaired phase-amplitude coupling (PAC) has previously been attributed to pTau in mouse models,^35^ and is altered in mild cognitive impairment and Alzheimer’s disease.^42^ Mid-gamma is associated with entorhinal inputs to CA1 superficial apical dendrites, detectable in CA1d SP at the peak of theta cycles in rodents.^43^ Despite neurodegeneration and pTau (Fig. 2f-h), higher amplitude mid-gamma oscillations were detected close to the theta peak (n=4/4 aged tg mice, 18-23 mo; Figs. 5e, S6b,c). Similar results were observed in aged ntg mice (n=3/3, 20-24 mo) and ageing tg mice (mean IMF frequency distributions and PAC; n=2, 13-14 mo). Theta frequency was typically lower during the peak-to-descending phase of theta oscillations (Figs. 5e, S6b,c).

Memory consolidation is associated with another form of cross frequency coupling: the sharp wave ripple (SWR) complex, consisting of 130-230 Hz ripple oscillations occurring during ~30-120 ms sharp waves.^44^ We detected ripples from CA1d SP in aged ntg and tg mice (18-24 mo) while they rested on the running disc (head-fixed) or in an open field. We found no difference between ntg (n=3) and tg (n=2) mice in terms of ripple frequency (124.7 ± 4.9 Hz ntg, 123.4 ± 4.6 Hz tg), ripple duration (53.1 ± 4.6 ms ntg, 51.7 ± 4.6 ms tg), or ripple cycle counts (6.6 ± 0.7ntg, 6.3 ± 0.4 tg). Similar results were found for ageing tg mice (n=2, 13-14 mo). This suggests that the sampled CA1d SWRs are unaffected by pTau. In summary, our analysis of network oscillations suggests unimpaired neuronal coordination in the hippocampus, consistent with good performance in hippocampal-dependent spatial working and reference memory tasks.

### Identification of single hippocampal neurons recorded in awake tg mice

After determining that local pTau did not affect network activity, we next investigated whether pTau disrupted the activity of individual neurons. To directly compare pTau+ and pTau− neurons, we performed extracellular recordings with glass electrodes in the hippocampal formation of head-restrained and freely-moving mice, followed by juxtacellular labeling (Fig. 6). We targeted the ProS, which contained high levels of pTau+ from an early age (Fig. 1). From a total of 47 recorded neurons (from n=8 aged tg mice, n=3 ageing tg mice and n=1 young tg mouse; 6-28 mo), we recovered 9 neurons (n=6 pyramidal neurons and n=3 interneurons). We tested 8 for AT8 (pTau) and all were immunonegative (Fig. 6d,e). We also recorded 20 single neurons in ntg littermates and wild type mice (from n=5 young mice, n=1 ageing mouse and n=1 aged mouse; 6-18 mo) and recovered 2 pyramidal neurons and 1 interneuron (Fig. 6a,b). Identified pyramidal cells in both tg and ntg mice were immunopositive for CTIP2 (Fig. 6b,e), matching the profile of pTau+ cells (Fig. 2a).

**Figure 6.**
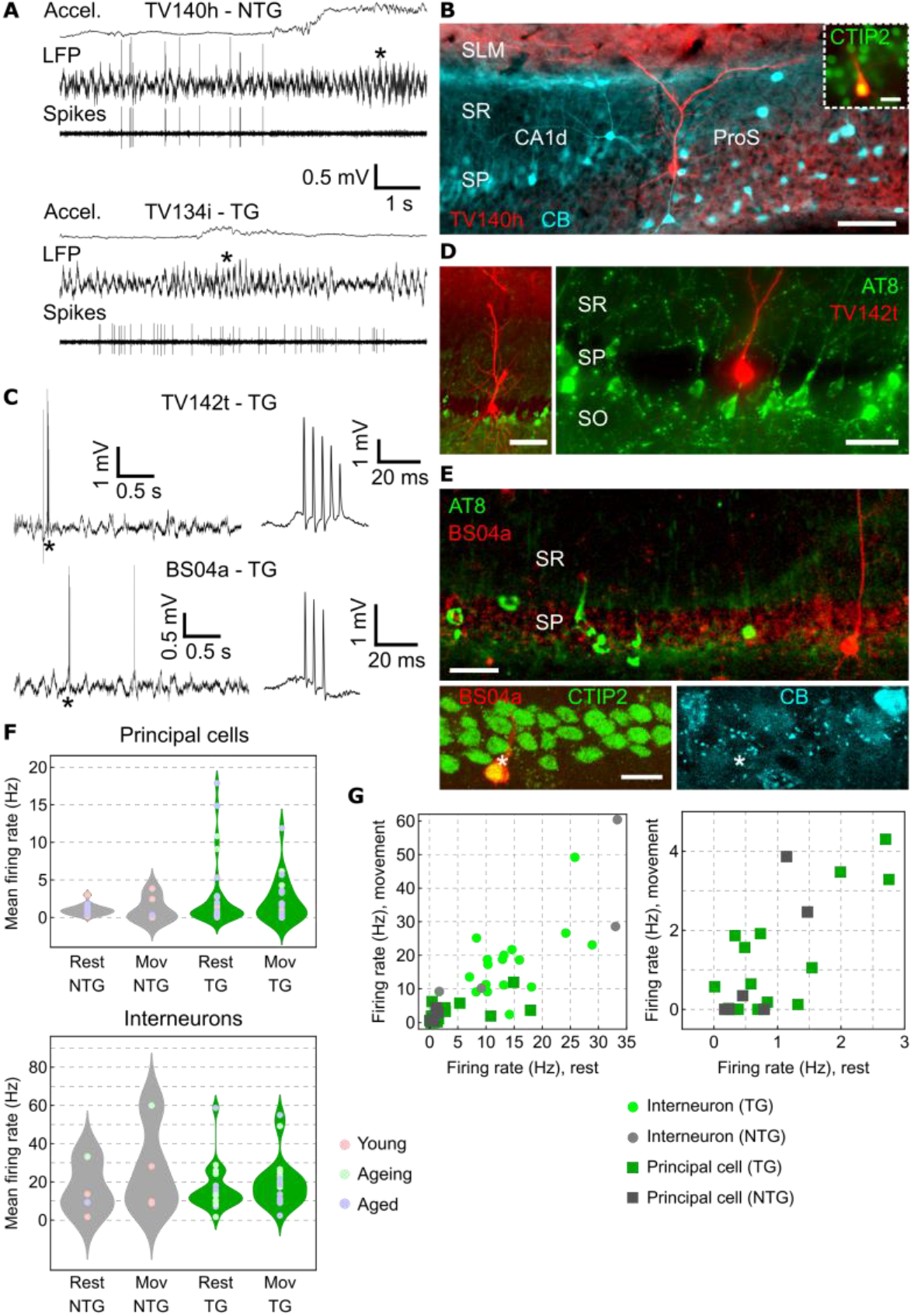
Identification of hippocampal neurons in relation to pTau. **(A)** Glass electrode extracellular recordings of pyramidal neurons in freely moving mice. Top, ProS LFP and associated band-pass filtered spikes from cell TV140h during spontaneous rest and exploration (18 mo ntg). Bottom, CA1d LFP (tungsten wire, contralateral) and spikes from cell TV134i (10 mo tg). Accel., acceleration. Asterisks, higher amplitude theta during movement. **(B)** Cell TV140h from A in ProS (neurobiotin, red) is CB– (cyan) and CTIP2+ (inset, green). Widefield epifluorescence. **(C)** Glass electrode recordings in head-fixed tg mice. Wideband LFP of large irregular activity and spikes during rest from cells TV142t (top, 18 mo) and BS04a (bottom, 13 mo). Right, CS at higher time resolution (asterisks). **(D)** Identified CA1 pyramidal cell TV142t (neurobiotin, red) from C lacks pTau (green). Widefield epifluorescence. **(E)** Identified CA1 pyramidal cell BS04a (neurobiotin, red) from C lacks pTau (green) and CB (cyan, asterisk) but is CTIP2+ (green, asterisk). Confocal maximum intensity z-projections (top, 48.5 μm thick; bottom, 9.25 μm thick). **(F)** Distribution charts of mean firing rates during rest and movement. Top, pyramidal neurons; bottom, interneurons; ntg, gray; tg, green. Points are individual neurons color coded by age group. **(G)** Firing rates for movement versus rest. Right, detail of lower firing cells from left. Scale bars (μm): B, D (left), 100; D (right), 50; E (top), 30; E (bottom), 15.

Under both freely-moving and head-fixed conditions, pyramidal neurons from both genotypes of all ages were recognized by relatively wide isolated spikes and complex spike (CS) bursts (Fig. 6a,c). Based on the locations, shapes and neurochemical profiles of the subset of recorded and labeled neurons (Fig. 6a-e) we could confidently classify unlabeled neurons with CS bursts as putative pyramidal cells, matching published data.^15,16^ Our sample of glass electrode-recorded pyramidal neurons had a typical mean firing rate distribution, consisting of a majority of low firing cells and few high firing cells,^45^ which broadly overlapped for the different age groups (Fig. 6f). Both genotypes had similar low firing rates (ntg rest, 0.92 [0.74] Hz, n=14 cells; tg rest, 0.69 [2.40] Hz, n=27 cells; *U*=191, *P*=0.9452; ntg movement, 0.17 [2.46] Hz, n=6 cells; tg movement 1.30 [3.45] Hz, n=22 cells; *U*=51, *P*=0.4132; Fig. 6f). Running speeds were also similar across genotype (medians of 6.14 versus 5.11 cm/s for ntg and tg mice, respectively; *U*=165, *P*=0.229).

Firing rates of labeled (identified) and unlabeled (putative) interneurons were typically higher than those of principal cells, lacked CS bursts, and showed a range of regular bursting activity and rhythmic firing patterns. Firing rates were also similar across genotypes (ntg rest, 11.53 [23.82] Hz, n=6 cells; tg rest, 13.17 [11.01] Hz, n=20 cells; *U* =58, *P*=0.9273; ntg movement, 18.95 [34.77] Hz, n=4 cells; tg movement 18.6 [12.05] Hz, n=19 cells; *U* =41, *P*=0.7765). Overall, the sampled range of firing rates of the recorded principal cells and interneurons in tg mice were similar to ntg littermates and other wild type (wt) mice (Fig. 6g).

### Fewer high-firing pyramidal neurons in aged transgenic mice during goal-directed navigation

Next, we investigated whether pTau affected neuronal activity when mice were engaged in a goal-directed navigation task. We trained water-restricted aged head-fixed mice to run on an air-suspended jet ball along a virtual linear track to reach a water reward. From 16 aged mice, of which 9 had previously been used in spatial memory tests, 11 could be successfully trained to run with a stable gait to the end of the corridor and initiate licking (n=6/9 tg and n=5/7 ntg mice; Table S1). Mice ran 45 ± 34 trials per session (1-4 sessions per mouse, with a maximum of 136 trials over 1 session for a tg mouse). For sessions with good coverage (>12 trials, 92% of all sessions), mice had similar trial durations (47 [64] s per trial across n=9 sessions for ntg mice versus 34 [30] s per trial across n=15 sessions for tg mice; *U*=41, *P*=0.1074). During this task we recorded multiple unit activity with acute silicon probes, unilaterally targeting ProS. We isolated 308 units (n=119 from 5 ntg mice, 16-24 mo; n=189 from 5 tg mice, 18-23 mo), localizing 147 units to ProS/SUB and 161 to adjacent CA1d (Table S1).

First we compared firing rates of all units that had complex spikes (CS) – putative pyramidal cells – matching the patterns of the recorded and labeled pyramidal neurons (Fig 6). There were no differences in mean firing rates between genotype (mean ± standard error 4.7±1.2 Hz for n=85 units from 5 ntg mice; 2.7±1.1 Hz for n=132 units from 5 tg mice; linear mixed model, mouse as a random factor, genotype fixed effect: *F*_(1,3.1)_=1.492, *P*=0.3072; Fig. 7a) or recording region (3.4±0.9 Hz for n=97 units from ProS/SUB; 4.0±1.0 Hz for n=120 units from CA1; region fixed effect: *F*_(1,13.6)_=0.378, *P*=0.5490). However, we found a significant difference for the interaction between genotype and region (5.5±1.3 Hz in ProS/SUB, n=39 units from 3 ntg mice; 1.3±4.0 Hz in ProS/SUB, n=58 units from 3 tg mice; 3.9±1.2 Hz in CA1, n=46 units from 4 ntg mice; 4.2±1.5 Hz in CA1, n=74 units from 3 tg mice; genotype-region fixed effect: *F*_(1,13.6)_=4.990, *P*=0.0428; Fig. 7b, Table S1). The distributions were also significantly different for ProS/SUB but not CA1 (pooled data per region, ProS/SUB ntg versus tg: *D*=0.3563, *P*=0.0037; CA1 ntg versus tg: *D*=0.1645, *P*=0.3751; Fig. 7b). These results suggest that there are fewer high-firing cells in ProS/SUB in aged tg mice.

**Figure 7.**
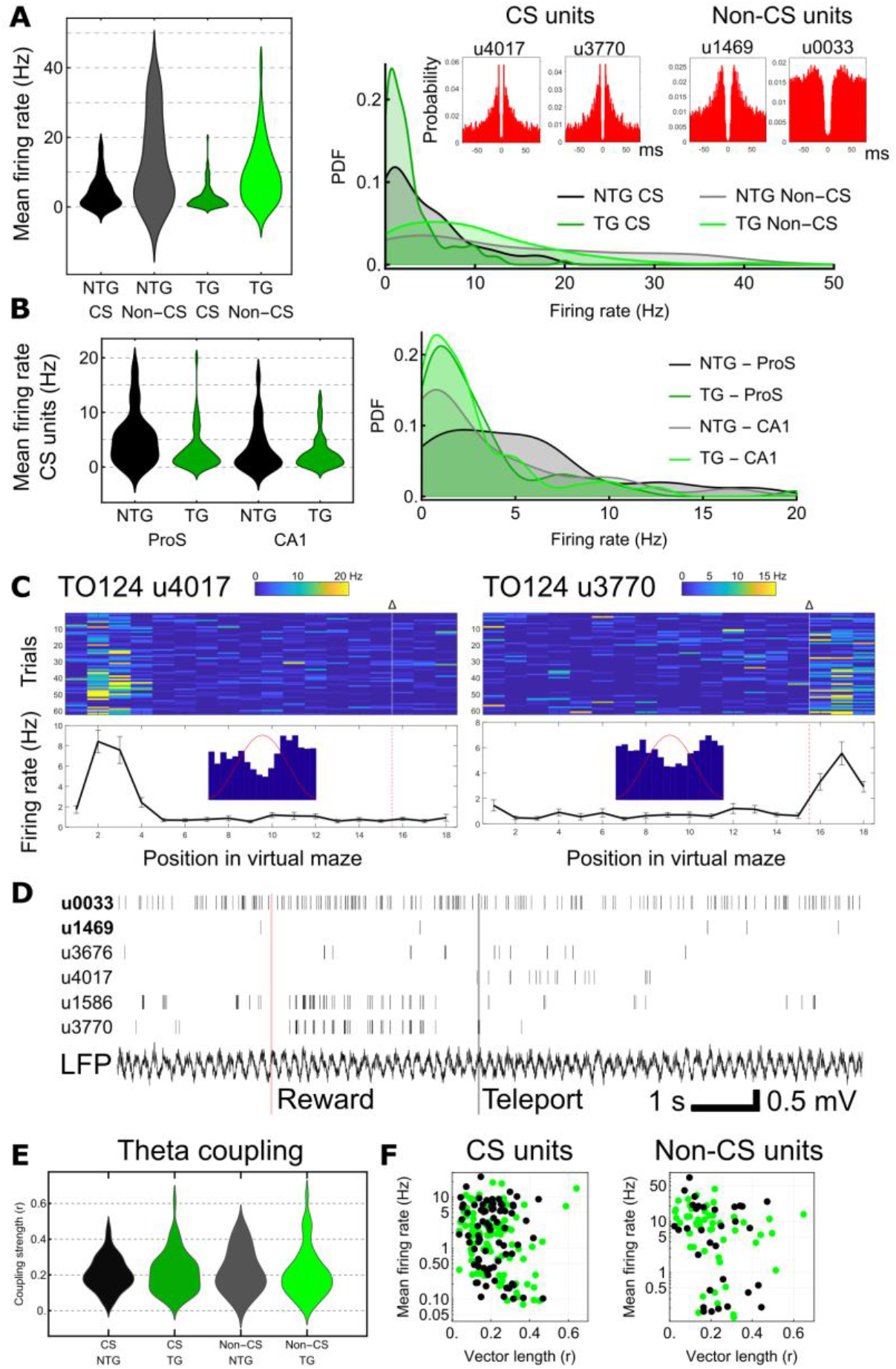
Loss of higher firing complex spiking cells but normal theta coupling in aged tg mice. **(A)** Left, distribution charts of complex spiking (CS) and non-CS units for ntg and tg mice (pooled data; one high-firing outlier removed from the ntg non-CS group). Right, probability density function (PDF). Red autocorrelograms feature units from a tg mouse (case TO124, 23 mo). **(B)** As in A but separated by region (ProS/SUB and CA1) for CS units only. **(C)** Two place cells from case TO124: unit (u) 4017 most active towards start of virtual linear track, whereas u3770 most active following reward (Δ) at end of track. Color bars, 1 to 99 % of linearized firing rate (blue min., red max.). Insets, theta phase histogram (20° bins; 180° corresponds to peak shown by red sinusoid). **(D)** Spike trains from 6 units (case TO124); non-CS units in bold. Note u1586 and u3770 are rhythmically synchronized after reward. ‘Teleport’, re-initialization of linear track (new trial). Also note coupling to LFP theta cycles, and spikes corresponding to place field of u4017 from C. **(E)** Distribution chart of coupling strength (vector length, r) for all units. **(F)** Firing rate during theta versus vector length for individual units. Black, ntg mice; green, tg mice.

Previous studies have set firing rate thresholds to help separate putative pyramidal cells (assumed to be low-firing) from high-firing interneurons.^46^ When we compared low-firing ntg and tg CA1 CS units (mean firing rates < 3 Hz) to a separate dataset obtained from younger wt mice performing a similar task on the same jet ball,^43^ we found no difference in firing rates (mean ± standard error 0.9±0.2 Hz, n=29 ntg units; 1.1±0.2 Hz, n=55 tg units; 1.1±0.1 Hz, n=159 wt units; linear mixed model, mouse as a random factor, genotype fixed effect: *F*_(2,9.6)_=0.6835, *P*=0.5278; Table S1, Figs. 7c,d, S6d). This suggests that for the subpopulation of low-firing pyramidal cells, aged mice have similar firing rate distributions to younger mice. Since we have identified high-firing (> 3 Hz) CS cells as pyramidal cells in ProS, SUB and CA1 (Fig. 6), we suggest higher firing ProS/SUBd pyramidal cells, which could have special roles distinct from other cells,^45^ are preferentially lost in tg mice, leaving the lower firing CS cells.

We also compared firing rates of units that lacked CS – putative interneurons – matching the profiles of some of the recorded and labeled interneurons. In contrast to CS cells, there were no significant differences in firing rates across genotype, region or their interaction (mean ± standard error 16.1±4.3 Hz for n=29 units from 5 ntg mice; 9.1±3.8 Hz for n=62 units from 5 tg mice; linear mixed model, mouse as a random factor, genotype fixed effect: *F*_(1,7.5)_=1.530, *P*=0.2532; 12.6±3.3 Hz for n=50 units from ProS/SUB; 12.6±3.4 Hz for n=41 units from CA1; region fixed effect: *F*_(1,42.8)_=0.0003, *P*=0.9874; 14.0±4.9 Hz in ProS/SUB, n=12 units from 3 ntg mice; 11.2±4.4 Hz in ProS/SUB, n=38 units from 3 tg mice; 18.3±4.6 Hz in CA1, n=17 units from 4 ntg mice; 6.9±4.9 Hz in CA1, n=24 units from 3 tg mice; genotype-region fixed effect: *F*_(1,42.8)_=1.534, *P*=0.2222; Fig. 7a, Table S1). The proportions of CS versus non-CS units were also similar across genotype (mean ± s.d. 76±17% CS units, n=5 sessions for ntg mice; 62±18% CS units, n=5 sessions for tg mice; excluding sessions with <14 units; *U*=18, *P*=0.2963; Fig. 7a, Table S1).

We also investigated whether reduction of high-firing ProS/SUB CS units in tg mice affected the relative proportion of place cells (Fig. 7c,d), which represent different locations of the virtual linear track.^47^ Ntg and tg mice had fairly similar proportions of place cells (17.5% (20/114 units), n=5 ntg mice; 25.3% (49/194 units), n=5 tg mice). Place fields were observed at different locations along the virtual corridor, with some associated with the reward (goal location, Fig. 7c,d). Interestingly, some units showed synchronized firing following reward periods (e.g. units u1586 and u3770, Fig. 7d). These data suggest that place cell mapping is preserved in these aged tg mice.

### Preservation of theta rhythmicity in aged mice

Given that we did not detect any major differences in theta oscillations or other associated spectral components (Fig. 5), we hypothesized that the remaining neurons in aged tg mice would retain their theta coupling, despite extensive pTau. Indeed, units from tg mice were coupled to theta and could become synchronized on individual theta cycles within overlapping place fields (Figs. 7d,e, S6d-f). For CS units, 95% from ntg mice (n=80/84 units) and 86% from tg mice (n=107/125 units) were significantly theta coupled (Rayleigh tests, *P*<0.05; Fig. 7c-f). There were no differences in theta coupling across genotype or region (mean ± standard error vector length r=0.21±0.013 for n=80 units from 5 ntg mice; r=0.22±0.011 for n=107 units from 4 tg mice; linear mixed model, mouse as a random factor, genotype fixed effect: *F*_(1,1.1)_=0.580, *P*=0.580; r=0.15±0.032 for n=59 units from ProS/SUB; r=0.24±0.024 for n=128 units from CA1; region fixed effect: *F*_(1,1.4)_=0.239, *P*=0.690; Fig. 7e). The most strongly coupled CS units for ntg and tg mice were r=0.47 and r=0.64, respectively; both were found within SUBd. We also found no difference in the distributions with respect to theta coupling versus firing rate (*D*=0.4458, *P*=0.0994; Fig. 7f). The phase distribution (spread) was greater in aged tg mice (circular standard deviation of angles, ProS/SUB: 40° for ntg mice (n=25 units) versus 71° for tg mice (n=34 units); CA1: 61° for ntg mice (n=55 units) versus 100° for tg mice (n=72 units); Fig. S6e). Furthermore, for the subsets of theta-coupled CS units with firing rates <3 Hz and significant phase coupling, coupling strength was no different from data obtained from younger wt mice ^43^ (mean ± standard error vector length r=0.21±0.015 for n=32 units from 4 ntg mice; r=0.23±0.011 for n=57 units from 3 tg mice; r=0.20±0.010 for n=152 units from 6 wt mice; linear mixed model, mouse as a random factor, genotype fixed effect: *F*_(2,*7.7*)_=0.988, *P*=0.415).

For non-CS cells, 94% from ntg mice (32/34 units) and 85% from tg mice (50/59 units) were significantly coupled to theta oscillations (Rayleigh tests, *P*<0.05). As with CS units, there were no differences in coupling strength of non-CS units across genotype (mean ± standard error vector length r=0.19±0.034 for n=32 units from 5 ntg mice; r=0.19±0.027 for n=50 units from 5 tg mice; linear mixed model, mouse as a random factor, genotype fixed effect: *F*_(1,6.2)_<0.001, *P*=0.987; Fig. 7e) or in the distributions of coupling versus firing rate (*D*=1.1272, *P*=0.6191; Fig. 7f). However, we detected significantly different coupling in ProS/SUB versus CA1 (r=0.15±0.032 for n=29 units from ProS/SUB; r=0.24±0.024 for n=53 units from CA1; region fixed effect: *F*_(1,13.0)_=5.524, *P*=0.035), suggesting that interneurons in CA1 are more strongly coupled overall. As for CS units, the phase distribution of non-CS units was also greater in aged tg mice (ProS/SUB: 49° for ntg mice (n=9 units) versus 82° for tg mice (n=20 units); CA1: 44° for ntg mice (n=23 units) versus 71° for tg mice (n=30 units); Fig. S6f). In summary, we could detect no differences in theta phase coupling in aged ntg and tg mice.

## Discussion

We found an age-related progression of pTau, consistent with previous reports,^9,13,14,48^ and other Thy1 promoter-driven 4R Tau lines.^49–51^ We further mapped pTau up to 28 mo, finding that deep-layer Wfs1+/CB– hippocampal pyramidal neurons preferentially accumulated pTau, followed by neurodegeneration. The apparent downregulation of the transmembrane glycoprotein Wfs1 in pTau+ pyramidal cells could be related to its role in regulating degradation of misfolded proteins in the endoplasmic reticulum, highlighting susceptibility to neurodegeneration.^18^ Oligodendrocytes also became pTau+ and were associated with pTau+ white matter tracts. We suggest the extensive Tau pathology in the hippocampal formation and loss of higher firing ProS cells contributed to specific deficits in short-term familiarity. We did not detect impairments in network oscillations or theta coupling, and spatial working and reference memory performance were similar for tg and ntg mice.

### Mechanisms and consequences of pTau spreading to oligodendrocytes

The pTau+ coiled bodies we found in oligodendrocytes resemble those in PSP, CBD and Alzheimer’s disease,^20–22,52^ and are probably a feature specific to overproduction of 4R pTau.^53^ Coiled bodies may facilitate pTau propagation in humans, as mice inoculated with pTau from tauopathy model mice or patients exhibit spreading via oligodendrocytes.^54–56^ This makes THY-Tau22 mice a useful model for aspects of PSP and CBD. As Thy1 is neuron-specific,^25^ we suggest pTau within axons spreads directly to oligodendrocytes. Alternatively, Thy1 in oligodendrocytes could be repressed in younger animals; examining aged Thy-GFP mice for GFP+ oligodendrocytes may be informative. Oligodendrocytes containing pTau along fiber tracts strongly suggests entry at paranodal septate-like junctions between the axonal membrane and myelin adjacent to the nodes of Ranvier,^57^ which could then spread to the cell body.^58^ In the *MAPT*^P301S^ tauopathy model, new oligodendrocytes increase at 5-6 mo prior to onset of cognitive deficits, which has been suggested to compensate for an early loss of myelin.^59^ Disruption of oligodendrocyte precursor cells causes memory impairments.^60^ Conversely, promotion of oligodendrogenesis in aged mice is associated with memory improvement.^61^ In THY-Tau22 mice, it is possible that disruption of myelin production affects action potential propagation. Myelinating satellite oligodendrocytes ^24^ are tightly associated with neuronal somata, and are electrically coupled to astrocytes and other oligodendrocytes.^62^ As we have observed coiled bodies close to pTau+ pyramidal neurons, and fewer higher firing pyramidal cells in aged tg mice, disruption of gray matter oligodendrocytes could also contribute to the reduction in higher firing activity and affect spike initiation and/or propagation. Anterograde transneuronal pTau spread is unlikely as we did not observe pTau in regions innervated by abundant pTau+ ProS/SUBd/CA1d pyramidal neurons such as lateral septum, anterior thalamus or mammillary bodies. However, ProS projects to the amygdala,^11^ and both regions were highly enriched in pTau.

### Unimpaired cortical neuronal coordination and spatial memory

Theta-coupled bursting GABAergic neurons of the medial septum (MS) project to the temporal cortex,^63–65^ and MS inactivation reduces cortical theta power and impairs spatial memory.^36,66–69^ Disruption of MS cholinergic neurons by pTau can also cause memory deficits,^70^ and a cholinergic deficit was reported in THY-Tau22 mice.^27^ Selective inactivation of rat anterior or ventral midline thalamic nuclei also results in spatial memory impairments;^71,72^ several of these nuclei also contain theta-rhythmic neurons and project to temporal cortical areas. We did not detect pTau+ neurons in the MS or thalamus, which could explain why theta coupled neurons, theta peak frequencies, IMFs, and cross-frequency coupling in heavily pTau-dominated CA1-SUB had broadly similar distributions for both genotypes under our conditions. We used longitudinal behavioral testing at different ages. This may have acted as an enriched environment, which is known to delay onset of memory deficits.^73^ Also, spatial learning can lead to neuroprotective oligodendrogliogenesis.^60^ Many behaviorally-tested animals were used for the neurophysiological recordings. Previous studies reporting that 9-10 mo tg mice spend longer finding the hidden platform in a Morris water maze also found they had increased thigmotaxis,^13,28^ with deficits emerging as early as 3 mo.^9^ We minimized this potential confound by using a water escape Y-maze,^29^ revealing tg mice are unimpaired for spatial reference memory. In summary, unimpaired hippocampal neuronal coordination may largely explain why we observed normal hippocampal-dependent spatial reference and working memory performance.

### Reduction in short-term familiarity

We found that >13 mo tg mice failed to discriminate novel and other arms of the hippocampus-dependent ^74^ spontaneous spatial novelty preference Y-maze, as shown previously in younger mice.^27,28^ This suggests impairment in recognizing familiarity of extra-maze spatial cues, which are required for spatial novelty preference.^75^ Encoding and recall of spatial memories involves coordination of sequential neuronal activity within and between different brain regions. Compared to CA1, the SUBd contains a richer variety of spatially-modulated neurons including head direction cells, boundary vector cells, grid cells, as well as place cells,^72,76^ and unlike CA1, receives direct glutamatergic input from the anterior thalamus. The loss of higher firing neurons in ProS/SUBd, due to high pTau levels, suggests sequences of neurons encoding different features of the environment may be disrupted. A reduction in short-term familiarity could also explain why tg mice spent significantly longer in the start arm of the T-maze before initiating runs: placement in the start arm retained some novelty for the mouse, possibly due to the reduction in ProS/SUBd activity. Lastly, we found that both young and ageing tg mice naïve to the elevated plus maze spent longer in the open arms. The ventral hippocampus and inter-connected amygdala also develop early Tau pathology,^14^ which we observed throughout the lifetime of the tg mice. Furthermore, lesions of rat ventral hippocampus alters anxiety-related contextual behavior but not spatial learning.^77,78^ Taken together, changes in spatial familiarity may be explained by potential deficits of different kinds of spatially-modulated cells within pTau-enriched ProS/SUBd.

### Effects of pTau on identified neurons

Our single recorded and labeled neurons lacked pTau, hence we do not know how pTau affects *in vivo* firing, but in aged tg mice there were fewer higher firing ProS pyramidal cells during goal-directed navigation. In addition to neurodegeneration in aged mice and potential disruption of axon potential propagation by pTau+ oligodendrocytes, it is also possible that local pTau accumulation within pyramidal neurons causes a reduction in firing rate, contributing to the shift in the distribution to lower rates. To our knowledge, there are no reports on *in vivo* firing of single hippocampal neurons that specifically accumulated pTau. In the rTg4510 tauopathy model, mice have slower membrane potential oscillations and reduced firing rates, but only one labeled pTau+ neocortical neuron was reported.^37^ *In vivo* calcium imaging in 6-12 mo rTg4510 mice show diminished activity under anesthesia.^79^ These lower activity levels could be attributed to an actual loss of higher firing cells, shifting the populations towards lower firing cells. Matching pTau content to imaged/recorded neurons is needed to test the effects of pTau on neuronal activity. In 3 male 3xTg mice, hippocampal place cells had reduced firing rates, lower place field stability and less spatial information compared to wt mice, which was attributed to abnormal theta and gamma oscillations.^80^ We did not record multiple place cells under freely moving conditions so do not know whether THY-Tau22 mice also exhibit similar deficits, although mutant Tau in combination with induced amyloid-beta plaques and potential extra-hippocampal transgene expression may have contributed to deficits in the 3xTg model.

Skewed firing rate distributions are a common feature of the brain, with the high-firing minority having key roles, tending to be more highly connected, and able to generalize across different contexts.^45,81^ A loss of strongly-potentiated higher firing cells would presumably result in an overall shift of the firing rate distributions towards a lower median. In acute slices, long term potentiation in CA1 was similar in tg and ntg mice, despite lower synaptic excitability in ageing tg mice.^9^ Chronic high-density recordings in ageing mice would be required to determine whether synaptic plasticity in lower firing cells largely compensates for the loss of highly potentiated synapses. The effect on spatial navigation could be that mice take longer to recognize contexts due to a loss of ‘generalizer’ high-firing neurons. Nevertheless, spatial memory *per se* is retained due to unimpaired neuronal coordination (such as spike timing during theta oscillations). Alzheimer’s disease, FTDP, PSP and other tauopathies show Tau pathology in several subcortical areas such as the thalamus and brainstem, which could promote memory deficits. Overall, the results of our detailed characterization of the THY-Tau22 mouse line demonstrate that in spite of widespread hippocampal Tau pathology, uptake of pTau by oligodendrocytes, neurodegeneration, and loss of high firing ProS/SUBd cells, many measured functional parameters and behaviors remain remarkably similar to age-matched control animals, likely due to the operations of unaffected neuronal systems.

### Limitations of the study

Since we recorded mostly in CA1d and ProS/SUBd, we cannot rule out changes in network oscillations and neuronal activity in other pTau-rich regions.^82^ We recorded place cells in head-fixed mice running along a virtual linear track. Establishing whether other kinds of spatially modulated cells are affected by pTau requires high density recordings in ProS/SUBd of freely moving aged tg mice. To directly compare pTau+ and pTau– neurons, *in vivo* calcium imaging experiments could be performed followed by *post-hoc* immunohistochemical tests for pTau matched to the imaged neurons, as we carried out for single recorded and juxtacellularly labeled cells.

## Acknowledgements

This study was supported by the Medical Research Council (MR/R011567/1), Alzheimer’s Society (522 AS-PhD-19a-010), Wellcome Trust (108726/Z/15/Z), Erasmus+, and the Nuffield Benefaction for Medicine and the Wellcome Institutional Strategic Support Fund (0007268). We thank Luc Buée and INSERM for kindly providing THY-Tau22 mice, Vitor Lopes dos Santos for advice on EMD, Balint Lasztoczi for published recording data, Emily Hunter and Verena Gautsch for assistance with histology, and Sawa Horie and Katalin Lengyel for providing training to B.S. for electron microscopy. We thank Alex Jeans for commenting on an earlier version of the manuscript.

## Author contributions

Conceptualization: T.J.V., B.S., A.T.O., K.H., J.S., D.B., P.S.; Methodology: T.J.V., B.S., A.T.O., K.H., J.S., D.B., P.S.; Investigation: T.J.V., B.S., A.T.O., K.H., J.S., D.B., P.S.; Writing – Original Draft: T.J.V.; Writing – Review & Editing: T.J.V., B.S., A.T.O., K.H., J.S., D.B., P.S.; Funding Acquisition: T.J.V., P.S.; Supervision: T.J.V., A.T.O., P.S.

## Methods

### RESOURCE AVAILABILITY

#### Lead Contact

Further information and requests for resources and reagents should be directed to and will be fulfilled by the lead contact, Tim Viney.

#### Materials Availability

This study did not generate new unique reagents.

#### Data and code availability

- Raw imaging and electrophysiological data reported in this paper will be shared by the Lead Contact upon request.
- All original code will be made publicly available as of the date of publication.
- Any additional information required to reanalyze the data reported in this paper is available from the lead contact upon request.

### EXPERIMENTAL MODEL AND SUBJECT DETAILS

A total of 97 mice were used in this study (~50 % were female), comprising 51 heterozygous THY-Tau22 mice (tg), 43 non-transgenic littermates (ntg), and 3 C57Bl6j mice (wt). The THY-Tau22 transgenic mouse line was created by the group of Prof Luc Buée, with Thy1.2 driving expression of human *MAPT* with G272V and P301S mutations.^9^ All animal procedures were approved by the local Animal Welfare and Ethical Review Body under approved personal and project licenses in accordance with the Animals (Scientific Procedures) Act, 1986 (UK) and associated regulations. We bred mice on a C57Bl6j background, as in the original publication. The oldest mouse was nearly 28 mo at the scientific endpoint (case TO117, tg female, glass electrode recordings; see below). Almost all aged mice reached their scientific end point (100 % of female ntg mice (n=16/16), 91 % of female tg mice (n=19/22), 94 % of male ntg mice (n=17/18), 100 % of male tg mice (n=13/13)).

Body weight was routinely monitored. Young male tg mice had reduced weight compared to ntg mice (~8 % difference; 31.6 ± 2.3 g ntg mice (n=11) and 26.8 ± 0.45 g tg mice (n=5); F_(1,14)_ = 20.3, p < 0.001, one-way ANOVA; 3-5 mo; Fig. S2d), confirming previously published data ^9^. Aged tg mice also had reduced body weight (38.3 ± 3.2 g male ntg (n=7), 28.8 ± 2.6 g male tg (n=7), 37.1 ± 0.77 g female ntg (n=4), 28 ± 1.4 g female tg (n=6); two-way ANOVA *F*_(1,20)_ = 67.4, *P* < 0.001 for genotype, *P* = 0.7680 for sex, *P* = 0.7467 for genotype-sex interaction; 18-21 mo; Fig. S2d). In even older (22-25 mo) mice, there was a significant weight difference between sexes (37.5 ± 2.5 g male ntg (n=6), 27.3 ± 1.5 g male tg (n=6), 33.1 ± 6.9 g female ntg (n=8), 24.8 ± 2.9 h female tg (n=11); *F*_(1,27)_ = 38.1, *P* < 0.001 for genotype, *P* = 0.0323 for sex, *P* = 0.5460 for genotype-sex interaction; Fig. S2d).

Of the 55 mice that underwent behavioral testing, 17 were subsequently implanted for electrophysiological recordings. A further 23 mice not tested behaviorally were also used for electrophysiological recordings. From this group, 3 aged tg mice (2 males, 1 female) had minor gait problems and tremor. Most mice were perfusion-fixed at their scientific endpoint, with 10/97 mice used only for histology. The remaining 8 mice were monitored for weight only and not used in procedures. Animals were maintained in individually ventilated cages with their littermates on a 12/12 h light-dark cycle (lights on at 07:00) and all behavioral tests were carried out during the daytime. Implanted mice were also grouped housed wherever possible. Room temperature was maintained at 21°C and humidity was between 50-60 %. Running wheels were placed in the cages of mice undergoing habituation/training to the jet ball (see below). All mice were fed on a standard RM3 diet (pellets, product 801700, Special Diets Services). Each animal was assigned a name (two letters followed by a code, e.g. **TT**21D, **TV**140), glass electrode recorded cells were assigned a lower case letter (e.g. TV140**h**, Fig. 6a), and units isolated from silicon probes were assigned a code (e.g. TO124 u4017, Fig. 7c).

### METHOD DETAILS

Electrophysiological recordings, behavioral tests, and cell counts were made blind to genotype.

#### Surgical procedures

Mice were anesthetized with isoflurane and the scalp was clipped. After subcutaneous injection of Buprenorphine (0.1 mg/kg dose, Vetergesic, Ceva), mice were fixed to a stereotaxic frame (Kopf Instruments) via ear bars and a jaw bar and maintained with 1–3 % (v/v) isoflurane on a homeothermic blanket (Harvard Apparatus). Ocular lubricant was applied to the eyes, and Bupivacaine (Marcaine, Aspen) was injected into the scalp. Under aseptic conditions, an incision was made along the scalp to expose the skull. Two M1 screws (Precision Technology Supplies) were fixed into the skull above the cerebellum, with one or both used as electrical reference. Another screw was fixed ~1.50 mm anterior of the bregma. After scoring the skull and applying a thin layer of cyanoacrylate super glue, a machined glass-reinforced plastic D-shaped head plate (custom made at the Department of Physics, Oxford University) was positioned over the screws and Refobacin bone cement (Zimmer Biomet) was used to secure the head plate and screws to the skull. A 0.7 g head plate was used for *head-fixed* and *freely moving mice*, and a slightly longer 1.1 g head plate was used for *jet ball* experiments. Mice that failed to learn for jet ball experiments could be used in head-fixed recordings. Similarly, if freely moving mouse recordings failed, head-fixed recordings could be carried out instead. For the experiments involving freely moving mice, a 7 stranded PerFluoroAlkoxy-coated steel wire (0.002 inch thickness, A-M Systems) was stereotaxically inserted into CA1d to target stratum pyramidale (−2.30 mm antero-posterior (AP) and 1.50 mm medio-lateral (ML) from bregma, 1 mm below the dura mater) and secured with bone cement and/or blue-light curing dental cement (Tetric EvoFlow, Ivoclar Vivadent). A custom-made stainless steel NanoMotor holder (Department of Pharmacology, Oxford University) was secured with bone cement to one side of the head plate. A headstage connector (a double row of pins) was also secured to the head plate, and the tungsten wire and reference were soldered to specific pins of the connector. Mice were administered a subcutaneous injection of 0.5 ml 5 % (w/v) glucose in saline (Aqupharm 3, Animalcare) peri-operatively. Craniotomy sites were as follows (mm from bregma): −2.90 AP, 1.75 ML (distal CA1d / proximal SUBd, also known as prosubiculum ^11,12^, for the majority of mice); −3.20 AP, 1.90 ML (SUBd); −2.50 AP, 1.70 ML, 10° posterio-anterior angle (CA1d). In some mice, craniotomies were performed during the first surgery immediately after head plate implantation. In all others (including all mice for jet ball experiments), craniotomy sites were marked by partially drilling into the skull, silicon (Smooth-On) was used to cover the skull inside the head plate, and mice were left to recover in a home cage over a heated blanket. After habituation/training (below), a second brief surgery was carried out, with the same anesthesia regime as above. Craniotomies were performed at the marked sites, and the dura was removed with a bent 27 gauge needle. Silicon was used to protect craniotomy sites and mice were left to recover in a home cage over a heated blanket. Electrophysiology experiments were initiated the following day.

#### Electrophysiology

##### Glass electrode recordings in head-fixed mice

Implanted mice were habituated to head-fixation using a custom-made stainless steel block (Department of Physics, Oxford University), secured to a heavy-duty frame (model 1430, Kopf Instruments). Mice could spontaneously run and rest on a 30 cm diameter plastic Frisbee (‘running disc’) covered with paper roll. On recording days, the silicon was removed and replaced with sterile saline. Two separate glass electrodes filled with 3 % neurobiotin (w/v) in 0.5 M NaCl (8–18 MΩ) were advanced into the craniotomy sites using IVM-1000 micromanipulators (Scientifica Ltd) to reach the hippocampus/subiculum bilaterally. Signals were amplified x1000 (ELC-01MX, BF-48DGX and DPA-2FS modules; npi Electronic GmbH, Tamm, Germany). Both wide-band (0.3 Hz to 8 kHz) and band-pass-filtered (action potentials, 0.8–8 kHz) signals were acquired in parallel for each glass electrode and digitized at 20 kHz (Power1401; Cambridge Electronic Design Ltd, Cambridge, UK). HumBugs (Digitimer Ltd) were used to remove 50 Hz noise. A video camera was used to monitor behavior. Speed was measured using a rotary encoder (HEDM-5500#B13, Avago Technologies) attached to the underside of the Frisbee. Data were recorded using Spike2 software (Cambridge Electronic Design). Local field potential (LFP) measurements and extracellular recordings of single neurons were made bilaterally during spontaneous movement and rest periods. Cells of interest were juxtacellularly labeled with 200 ms positive current pulses, followed by a recovery period of at least 4 h.

##### Glass electrode recordings in freely moving mice

Mice were head-fixed as above and the craniotomy site was exposed (the contralateral site had the implanted tungsten wire). A custom-made piezo-electric NanoMotor (Kleindiek Nanotechnik and npi Electronic) was secured in the holder on the head plate and a miniature preamplifier/headstage (custom-made by npi Electronic) was attached to the headstage connector. A glass electrode containing 3 % neurobiotin in 0.5 M NaCl was attached to the NanoMotor and lowered into the brain. The mouse was released from head restraint and placed in a 50 cm^2^ open field with 30 cm high walls. Occasionally objects were added to the environment. Acceleration was measured with a built-in accelerometer and motion was tracked with a video camera and integrated LEDs on the headstage or by body-motion capture. Extracellular single neuron recordings and LFP measurements were made as for head-fixed mice, followed by juxtacellular labeling. Mice were returned to head-fixation in order to remove the NanoMotor and miniature headstage or continue head-fixed recordings as above.

##### Jet ball recordings

Mice were first handled for at least 2 d then *ad libitum* access to water was removed to initiate water restriction. Mice were gradually habituated to head-fixation over an air-suspended Styrofoam jet ball (part of a virtual reality setup, PhenoSys GmbH, Berlin, Germany). Mice were trained to run on the jet ball to advance a virtual linear track (projected in a dome). Once they reached the end, licking would release a 10 μl water reward (defining the end of a trial). Animals were teleported to the start of the virtual corridor to initiate the next run. Running followed by licking at the reward location was defined as goal-directed navigation. Each animal was trained twice daily for ~30 min for 8-20 d. Throughout the training days, the length of the virtual corridor was increased stepwise to make sure animals were motivated to run to receive the reward. The reward tube was adjusted before each session to ensure animals could reach the water droplets upon delivery. Mice were closely monitored to receive at least 1 mL of water each day to avoid the weight dropping below 10 % of their initial body weight. At least one day per week animals received *ad libitum* water; food was always available. The day after the craniotomy, and on subsequent recording days, an acute 16-channel silicon probe (Buzsaki16-A16 probe, A1×16 linear probe or a A2×2 tetrode probe, NeuroNexus) was painted with DiI or DiO, connected to an RA16-AC preamplifier (Tucker-Davis Technologies) and gradually lowered into the hippocampal formation (Table S1). Up to 4 recording sessions were carried out per day (one hemisphere per session). The hippocampus was identified based on approximate depth, high power theta oscillations during running, sharp waves during rest, polarity of theta and sharp waves across the different channels, and the presence of high firing neurons in an ~100 μm band corresponding to stratum pyramidale. Signals were amplified x1000 (Lynx-8 amplifiers, Neuralynx), band-pass filtered (0.5 Hz to 8 kHz), digitized at 20 kHz with a Power1401 and recorded with Spike2. Jet ball movement, position along the virtual track, a lick sensor and reward delivery were synchronized to the recordings. A video camera was used to monitored behavior. Four implanted mice failed to learn goal-directed navigation (cases: TV147, female, ntg, 23 mo; TV148, female, ntg, 23 mo; TV164, male, tg, 17 mo; TV166, female, tg 19 mo) due to poor hindlimb gait and/or tremor. Instead, they were used for head-fixed recordings as above. One other implanted mouse had poor balance (case TV169, female, tg, 17 mo) and could not be trained or used for head-fixed recordings so was excluded. Silicon probe recording sites were confirmed by examining DiI/DiO fluorescence in serially processed brain sections.

#### Data analysis

Data were analyzed in Mathematica (Wolfram Research), Python, MATLAB (The MathWorks), and Spike2.

##### Local field potentials (LFPs)

In Spike2, DC shifts were removed with 0.3 s sliding windows then channels were downsampled to 1.25 kHz. Channels were either exported for analysis with other programs, or analyzed further in Spike2. Theta epochs were defined by the power ratio of theta (5-12 Hz) over delta (2-4 Hz) in 2 s windows (root mean square (rms) amplitude) being greater than 2. To calculate the peak theta frequency, LFPs recorded with glass electrodes were sampled at different depths of CA1d, predominantly from stratum pyramidale. Up to 5 theta epochs were sampled per recording, ranging from 4 to 34 s to define each epoch. A Hanning window (FFT size 2048, 0.6 Hz resolution) was used to compute the power spectral density in Spike2. The theta peak frequency was measured directly from the curve (interpolated with a cubic spline). Values obtained from 2-4 recordings per mouse were binned and averaged for low running speeds (4-6 cm/s) and ‘high’ speeds (6-10 cm/s). We also measured the theta peak from 6 mice (3 ntg, 3 tg) running in virtual reality; 3 theta epochs were randomly sampled from 1 recording per mouse from a representative channel of an acute silicon probe targeted to distal CA1d. Theta epochs were between 10 and 218 s (mean of 94 ± 61 s). Actual speed could not be determined for these recordings. Ripples (band-pass filtered, 130-230 Hz) were detected from glass electrode LFP recordings by selecting periods where the rms amplitude (power) was greater than 3 standard deviations above the mean power. At least 2 recordings were sampled per animal, ranging from 31 to 94 detected ripples.

##### Hilbert-Huang transform.^83^

Empirical mode decomposition (EMD) was carried out in Python using the emd package (https://gitlab.com/emd-dev/emd) and its dependencies.^40^ Sampled glass electrode recordings ranged from 14 to 100 s of unfiltered LFP containing mostly movement periods. For silicon probe recordings, sampled LFPs containing running periods on the jet ball ranged from 57 to 182 s. To obtain intrinsic mode functions (IMFs), the following masks were used for masked EMD (emd.sift.mask_sift): (350, 200, 70, 40, 30, 7, 1) divided by the sampling rate. Instantaneous amplitude, frequency and phase were obtained for each IMF using emd.spectra.frequency_transform with the Normalized-Hilbert Transform (nht). We excluded IMF1 as it contained mostly noise, and excluded any mid-gamma range IMF that had 50 Hz electrical noise or poor movement periods (low signal to noise ratio). Consequently, 1 ntg and 1 tg mouse were excluded for IMF3. Phase amplitude coupling was analyzed using emd.spectra.hilberthuang, binning the theta IMF instantaneous phase values. We excluded 1 aged tg and 1 ntg mouse due to 50 Hz electrical noise. We also excluded 1 ntg mouse that lacked sufficient movement for PAC detection, and another ntg mouse due to the glass electrode positioned within stratum radiatum.

##### Single neuron glass electrode recordings

Recording locations were estimated from at least two of the following features: relative depth from a juxtacellularly-labelled recovered cell on the same pipette tract; LFP profile (e.g. positive sharp waves (deep layers) or negative going sharp waves (superficial layers), presence of ripples (highest amplitude in pyramidal cel layers), shape and amplitude of theta oscillations, cross frequency coupling of mid-gamma oscillations to the peak of theta cycles (CA1 stratum pyramidale), presence or absence of ‘dentate spikes’ for the dentate gyrus); evidence of a pipette tract and/or a cluster of weakly-labeled cells marking a recording location. Unlabeled/non-recovered prinicipal cells were compared to recovered (labeled) cells based on the presence or absence of complex spike bursts. Only data acquired before juxtacellular labeling were used for analysis. For firing rates, spikes within a complex spike (CS) burst were counted individually.

##### Acute silicon probe recordings

Spikes were detected, sorted, and clustered offline using Kilosort2.^84^ Subsequently, interactive visualization and manual curation of the data were carried out using phy software ^85^ in order to obtain well-isolated and stable units based on refractory periods, spike waveform and cross-correlations.^38^ Units with firing rates < 0.1 Hz were excluded from further analysis. The first trial from each session and any trials showing electrode drift were excluded. Data analyses were performed using built-in or custom-written software in MATLAB and Spike2. Firing rates (spikes/s) were computed across 18 sequential bins: We split each run into 15 spatial bins covering the virtual corridor followed by 3 temporal bins (1 s each) covering the end of the corridor corresponding to the reward location (the virtual environment was locked for 3 seconds when mice reached the end of the corridor in order to receive the water reward). Units were classified as complex spike (CS) cells if they showed a prominent peak between 3-8 ms in their spike time autocorrelograms followed by a relatively fast decay.^38,47^ Units were compared to a separate dataset from,^43^ who defined putative pyramidal cells (PCs, Table S1) as units with “an overall firing rate < 3 Hz, a spike width at 90% of the peak amplitude above 0.5 ms, and an event autocorrelogram value below 10 ms”. For firing rate analysis, one session each was excluded for animals TV149 and TO125 due to probes being located in the isocortex, just dorsal of the hippocampus (6 units in total), and one session was excluded for animal TV179 due to a lack of isolated units.

##### Place cell classification

Significant peak firing rates along the virtual corridor were computed by a permutation test as described by Ozdemir et al.^86^ Each unit’s firing rate was calculated across 18 spatio-temporal bins. Then, we randomly shuffled these time windows within a run for every trial, calculating a surrogate firing rate. Upon repeating this procedure 10,000 times, we obtained the *P* values of spatial firing modulation by comparing the actual firing rate to the distribution of surrogate rates. Subsequently, significance limits were corrected using a false discovery rate method for multiple comparisons. Well-isolated units were deemed place cells if they had a significantly higher firing rate in at least one spatial bin along the virtual corridor.

##### Theta coupling

A representative silicon probe site was selected for theta coupling analysis based on the profile of the unfiltered LFP, the extrapolated position from the recovered probe site (DiO or DiI tract), and the probe map. Mostly positive deflections in the LFP during non-running periods and typically high power theta oscillations during running indicated a likely stratum pyramidale or stratum oriens location. After detecting theta epochs as for glass electrode recordings above, theta phase was calculated by linear interpolation between troughs of the band-pass-filtered (5-12 Hz) theta oscillations, with 0° and 360° set as the troughs. The Rayleigh test was used to test for uniformity of circular phase distributions; the mean phase and mean vector length (r) were used as measures of the preferred phase and coupling strength, respectively (using custom scripts written in Mathematica). For ntg mice, we sampled 968-1,745 s duration theta periods (72,523 theta cycles) and excluded CS units and non-CS units that had fewer than 72 and 241 spikes, respectively. For tg mice, we sampled 291-3,023 s duration theta periods (91,233 theta cycles) and excluded CS cells and non-CS cells with fewer than 45 and 168 spikes, respectively.

#### Behavioral tasks

##### Spatial novelty preference Y-maze task

This task was designed to assess short-term spatial memory with the expectation that normal mice prefer novel over familiar spatial environments.^26,75^ A solid black plastic Y-maze (30 cm arms, 7 cm wide, 20 cm high walls) was positioned 17 cm from the floor. There were no intra-maze cues except for a 20 cm high magenta block that blocked one of the arms. Prominent extra-maze cues were used. One arm was designated as the ‘novel arm’, another the ‘start arm’, and the third as the ‘other arm’. The novel arm was blocked off and the mouse was placed in the start arm and was able to freely explore the start and other arms (exposure phase). After 5 minutes, the block was removed and mice could explore all 3 arms (test phase). Each mouse was only tested once, and arm assignment was counterbalanced for genotype and cohort. The maze was cleaned with ethanol before each test. Time spent in each arm was calculated manually or using ezTrack;^87^ total distance traveled (path length) was calculated using ezTrack. We excluded 3 mice from the path length analysis due to obstructions in the videos.

##### Water escape Y-maze task

This task was used to assess spatial reference memory without the confound of anxiety-related thigmotaxis, which is sometimes evident in the open field water maze.^29–31^ An enclosed Y-maze made of clear Perspex (30 cm arms, 8 cm wide, 20 cm high walls) was filled with water made opaque by mixing in white ‘ready mix paint’ (Reeves). A hidden inverted glass beaker (~6 cm diameter) was kept submerged at the end of one of the arms (‘goal arm’). Extra-maze cues were kept constant. At the start of each trial, the mouse was placed in one of the other two arms (‘start arms’). The start arm was allocated pseudorandomly across trials for each mouse with no more than 2 consecutive starts from the same arm and equal numbers of starts from the arm to the left and right of the target arm in each block of trials). The goal arm remained the same for each animal, but the allocation of the goal arm was counterbalanced for genotype and cohort. The mouse was allowed to swim freely until it reached the hidden platform. However, if it did not reach the platform by 90 s it was guided by the experimenter. The mouse was allowed to remain on the platform for 30 s before being removed. There were 5 trials/d for 10 days (acquisition training) with an inter-trial interval of 15 s. For each trial, the first arm entered (choice) was scored 1 for the goal arm and 0 for the other arm or re-entered the start arm. A block was defined as 5 trials, with performance expressed as percentage of correct first choices (choice accuracy). On day 11, a probe trial was performed, whereby the platform was removed and the mouse was allowed to swim freely for 60 s. One ntg mouse was excluded from analysis as it failed to make choices on days 4-10.

##### Non-matching to place T-maze task (rewarded alternation)

This task assessed spatial working memory.^32^ The T maze consisted of a 50 × 10 cm start arm and two identical 30 × 10 cm goal arms. Walls were 10 cm high, and the maze was raised 44 cm off the floor. Group-housed mice were maintained on a restricted feeding schedule to ~90 % of their free-feeding weight and fed every day at 16:00. On the 6^th^ day after starting food restriction, mice were habituated to 50% sweetened condensed milk in a dish in their home cage in the holding room. The next day, milk was provided in their cages inside the behavior-testing room (context 1). The following day, cagemates were group habituated to the T-maze with 0.5 ml milk rewards in plastic dishes at the ends of both goal arms. The room was well-lit and contained extra-maze cues. An odorless cleaning solution was used to clean the maze before testing different cages. After several more days on food restriction, non-matching to place testing began. Each trial consisted of a ‘forced run’ and a ‘choice run’, with 4 trials per day defined as a session. In the forced run, one goal arm was blocked with a wooden block. Mice were placed into a blocked-off start box at the far end of the start arm. After this start block was removed, mice had to run to the unblocked goal arm for a 0.1 ml reward. The blocked goal arm was determined by a pseudorandom sequence (with equal numbers of left and right turns per session, with no more than 2 consecutive turns in the same direction). If the mouse did not initiate a run within 60 s after removing the start black, the mouse would be re-tested after all the other cagemates were tested in that session. For all trials, the delay time in task initiation was measured from start block removal until leaving the start box area. After sampling the reward on the forced run, the mouse was removed from the maze. The mouse was placed back at the far end of the start arm and now had to choose between the goal arms (now with both arms unblocked). The average time between sampling the reward on the forced run to being free to initiate the choice run from the far end of the start arm was 16 s (from n=3,240 trials). The mouse was rewarded for choosing the unvisited (previously blocked) arm, i.e. for alternating. An alternation score was calculated by scoring 1 for alternating and 0 for not alternating. Mice were tested one trial at a time, so that all cagemates were tested once before starting their second trial (~4-8 min inter-trial interval, depending on cage size). A total of 10 sessions were undertaken for each mouse (40 trials total), and analyzed as blocks of 4 trials. The same mice were tested at 3 different ages, with the 2^nd^ and 3^rd^ tests no requiring habituation. Since ntg and tg mice reached high levels of performance (see Results), differences between genotypes might be revealed by using an unfamiliar context. In context 2, a different testing room was used featuring different extra-maze cues, an R-Carvone odor, and prominent intra-maze cues (white stripes on the maze floor and a red cross on the block), and cleaning was carried out with ethanol instead of the odorless cleaning solution.

##### Elevated plus maze

To test anxiety-related behavior, mice familiar to other behavioral tasks were placed individually in the center of an elevated plus maze facing an open arm. Two versions were used: one had 36 x 7 cm open and closed arms, 21.5 cm high gray wooden walls for the closed arms, and positioned 69 cm from the floor (for 3 mo mice). The other had 32 x 9 cm arms, 19.5 cm high black plastic walls for the closed arms, and was 92 cm from the floor (for 12 mo mice). The time spent in open and closed arms was measured over 5 min, excluding the junction.

#### Histology and imaging

##### Tissue processing

Mice were deeply anesthetized with sodium pentobarbital (50 mg/kg, i.p.) and transcardially perfused with saline followed by 4% paraformaldehyde, 15% v/v saturated picric acid, and 0.05% glutaraldehyde in 0.1 M phosphate buffer (PB), pH 7.4. Some brains were post-fixed overnight in fixative lacking the glutaraldehyde. After washing in 0.1 M PB, 70 μm coronal sections were cut using a Leica Microsystems VT 1000S vibratome and stored in 0.1 M PB with 0.05% sodium azide at 4°C. To visualize neurobiotin-labeled neuronal processes, tissue sections were permeabilized in Tris-buffered saline (0.9% NaCl buffered with 50 mM Tris, pH 7.4; TBS) with 0.3% Triton X-100 (TBS-Tx) or through rapid 2x freeze–thaw (FT) over liquid nitrogen (cryoprotected in 20% sucrose in 0.1 M PB) then streptavidin-conjugated Alexa Fluor 488 was applied at 1:1000 dilution in TBS-Tx (or TBS if permeabilized with FT) for 4 h at room temperature (RT) or overnight at 4°C. Sections were washed in TBS/TBS-Tx and mounted to slides in Vectashield (Vector Laboratories).

##### Immunohistochemistry

Sections were blocked for 1 h in 20 % normal horse serum (NHS) in TBS/TBS-Tx followed by 3 d incubation at 4°C in primary antibody solution containing 1 % NHS in TBS/TBS-Tx. The following primary antibodies (and dilutions) were used: Mouse anti-AT8 1:5000 (MN1020, lot ND 169248, Thermo Fisher Scientific; this antibody specifically recognizes Tau phosphorylated at serine 202 and threonine 205 residues of human Tau ^88^), goat anti-Calbindin 1:500 (Calbindin-Go-Af1040, Frontier Institute), rabbit anti-Calbindin 1:5000 (CB-38, lot No.5.5, Swant), rabbit anti-Calretinin 1:1000 (7699/3H, lot 18299, Swant), rabbit anti-Ctip2 1:1000 (25B6, ab18465, Abcam), rabbit anti-Olig2 1:500 (AB9610, lot 0603024913, Millipore), guinea pig anti-Parvalbumin 1:250 (195004/1-19, Synaptic Systems), rabbit anti-Parvalbumin (PV 27, lot 2014, Swant), rabbit anti-PCP4 1:1000 (sc-293258, lot GO814, Santa Cruz), goat-anti Satb1 1:250 (N-14, sc-5989, Santa Cruz), rabbit anti-Satb2 1:1000 (ab34735 and ab92446, Abcam), chicken anti-Tyrosine hydroxylase 1:500 (ab76442, Abcam), rabbit anti-Wfs1 1:1000 (11558-1-AP, lot 00043335, Proteintech). Specificity references can be found in the following publications: ^63,64,89,90^. Sections were washed 3 times in TBS/TBS-Tx then incubated in secondary antibody solution containing 1 % NHS in TBS/TBS-Tx for 4 h RT or overnight at 4°C. The following secondary antibodies (and dilutions) were used in various combinations (all raised in donkey): 1:250 anti-guinea pig and anti-rabbit DyLight 405 1:250 (706-475-148), anti-rabbit Violet421 1:100-200 (713-675-152, 705-605-147, 711-475-152), anti-mouse and anti-rabbit Alexa Fluor 488 1:1000 (711-605-152, 711-545-152), anti-rabbit and anti-chicken Cy3 1:400 (711-165-152, 703-165-155), anti-goat Cy5 1:250 (705-605-147), anti-goat Cy51:250 (705-605-147), anti-guinea pig and anti-rabbit Alexa Fluor 647 1:250 (706-475-148, 711-605-152) from Jackson ImmunoResearch and Alexa Fluor 405 (A31556) from Invitrogen. After 3 washes, sections were mounted in Vectashield.

##### Fluorescence microscopy

Sections were visualized and imaged with widefield epifluorescence using a Leitz DMRB microscope (Leica Microsystems) equipped with PL Fluotar objectives (magnification/numerical aperture: 5x/0.15, 10x/0.3, 20x/0.5, 40x/0.7; OpenLab software) or an AXIO Observer Z1 microscope (LSM 710; Zeiss) equipped with Plan-Apochromat 10x/0.3, 20x/0.8 and 40x/1.4 objectives (Axiovision or ZEN Blue 2.6 software). For confocal microscopy, the LSM 710 was used with Plan-Apochromat 40x/1.4, 63x/1.4, and 100x/1.46 objectives (ZEN 2008 5.0 or ZEN Black 14.0 software). Laser lines (solid-state 405 nm, argon 488 nm, HeNe 543 nm and HeNe 633nm) were configured with the appropriate beamsplitters. The pinhole was set to ~1 Airy Unit for each channel.

##### Fluorescence measurements and cell counts

To measure the intensity of AT8 immunoreactivity (Fig. 1b), 10x magnification epifluorescence tiles were made of the hippocampal formation. Three 300 μm^2^ regions of interest (ROIs) were selected, covering the ProS and two regions of CA1d. The mean pixel intensity was normalized relative to a background ROI. At least 2 mice per age group were measured. For quantification of CB and pTau colocalization, we examined pTau+ somata within the pyramidal cell layer, randomly sampling ~20 (range 7-28) pTau+ cells per case for 3-5 cases per age group. To count the number of CA1 pyramidal cells in age-matched ntg and tg mice (Fig. 2H), we imaged SATB2 immunopositive or DAPI stained nuclei with a 20x/0.5 lens focused on the top of 70 μm-thick coronal sections containing CA1d starting from proximal CA1 adjacent to the SATB2 immunonegative CA2 region. Counting was carried out using the Cell Counter plugin in Fiji (ImageJ). We averaged counts from 3 hemispheres per mouse, then compared the resulting mean counts from 2-3 mice per genotype. All quantifications were blind to genotype and age.

##### Single- and double-labeling pre-embedding immunohistochemistry and electron microscopy

Floating sections from cases TT21D (22.5 mo female) and TT33G (21.5 mo female) were washed 3x in TBS and blocked in 5% normal goat serum (NGS) in TBS for 45 min. Next, sections were incubated with AT8 and/or Olig2 primary antibodies in TBS for 6 d at 4°C. The following immunoreactions were performed: (1) One primary antibody with immunogold labelling followed by silver intensification; (2) one primary antibody with peroxidase reaction; (3) two primary antibodies, visualized by a silver-intensified immunogold reaction, followed by an immunoperoxidase reaction; (4) control, no primary antibody, immunogold and biotinylated secondary antibodies followed by silver-intensification. After incubation with primary antibodies, sections were rinsed three times in TBS and blocked for 30 min at RT in 0.1 % Cold Water Fish Skin Gelatin (CWFS) solution containing 0.8% NGS and 0.05% sodium azide in TBS to reduce non-specific background reactions. Next, sections were incubated overnight at 4°C in 1:300 biotinylated goat anti-rabbit IgG (Vector Labs, BA-1000) and 1:200 donkey anti-mouse ultrasmall immunogold (Aurion, 100.322) in CWFS solution. Sections were washed several times in TBS and once in 0.1 M PB followed by incubation with 2% glutaraldehyde in PB to fix the antibodies conjugated to immunogold particles. After repeated washes in TBS, sections were incubated overnight at 4°C in avidin-biotinylated horseradish peroxidase complex (Vectastain ABC Elite kit, Vector Laboratories) in TBS. To visualize immunogold particles, they were intensified with silver: first the sections were treated with enhancement conditioning solution (ECS; Aurion) diluted 10x with distilled water for 3×5 min. To enlarge the gold particles further, sections were then incubated in silver enhancement solution (SE-LM, Aurion) for 20 min at 20°C and washed in ECS. Subsequently, peroxidase was visualized using 3,3-diaminobenzidine (DAB; 0.5 mg/ml, Sigma-Aldrich) as chromogen developed with 0.01% H_2_O_2_. The same method and reagents were used for single and double labeling. After washing sections in PB, they were treated with 0.5% OsO_4_ (a fixative and heavy metal contrasting agent) in 0.1 M PB for 20 min on ice to control reaction time. Next, dehydration of sections was carried out in ascending ethanol series (50%, 70%, 90%, 95%, 100%) and in acetonitrile. To enhance contrast for electron microscopy, sections were incubated in 1 % uranyl acetate diluted in 70% ethanol for 25 min between the 70% and 90% ethanol steps. Subsequently, sections were placed in epoxy resin (Durpucan AMC, Fluka, Sigma-Aldrich), incubated overnight at RT and transferred onto slides. For polymerization, sections were incubated at 60°C for 2 d. Selected regions of proximal SUBd and/or distal CA1d were re-embedded into Durcupan blocks. A series of ~50 nm-thick sections were cut with an ultramicrotome (Leica Ultracut UTC) and collected on pioloform-coated copper grids. Finally, sections were counterstained with lead citrate and studied on a Jeol 1010 transmission electron microscope equipped with a digital GATAN Orius camera (Department of Physiology, Anatomy and Genetics, Oxford University).

### QUANTIFICATION AND STATISTICAL ANALYSIS

Mean and standard deviation (s.d.) are reported as mean ± s.d.; median and interquartile range (IQR) are reported as median [IQR]. Experimental units (e.g. mice, cells) are specified in the text after the n values. Statistical analysis was carried out in Mathematica, GraphPad Prism, and SPSS (versions 25 and 28, IBM). The alpha was set to 0.05. For data that approximated a normal distribution, unpaired Student’s T tests were used to compare two groups, otherwise Mann-Whitney tests were used. For comparisons of more than two groups we used Analysis of Variance (ANOVA) for parametric data. To account for variability in samples from different mice for silicon probe recording data (firing rates and vector lengths), we used a linear mixed model in SPSS with mouse as a random factor, and genotype, region and ‘cell type’ (CS, non-CS) as fixed factors. For analysis of circular data, we used the Rayleigh test (*Z*). The following test statistics are specified in the text and refer to the following tests: *D*, Kolmogorov-Smirnov; *U*, Mann-Whitney; *t*, unpaired T test; *F*, ANOVA.

---

| REAGENT or RESOURCE | SOURCE | IDENTIFIER |
| --- | --- | --- |
| Antibodies |  |  |
| AT8 | Thermo Fisher Scientific | MN1020, RRID: AB_223647 |
| Calbindin | Frontier Institute | Calbindin-Go-Af1040, RRID: AB_2571569 |
| Calbindin | Swant | CB-38, RRID: AB_10000340 |
| Calretinin | Swant | 7699/3H, RRID: AB_10000321 |
| Ctip2 | Abcam | ab18465, RRID: AB_2064130 |
| Olig2 | Millipore | AB9610, RRID: AB_570666 |
| Parvalbumin | Synaptic Systems | 195004/1-19, RRID: AB_2156476 |
| Parvalbumin | Swant | PV 27, RRID: AB_2631173 |
| PCP4 | Santa Cruz | GO814 |
| Satb1 | Santa Cruz | sc-5989, RRID: AB_2184337 |
| Satb2 | Abcam | ab34735, RRID: AB_2301417 |
| Satb2 | Abcam | ab92446, RRID: AB_10563678 |
| Tyrosine hydroxylase | Abcam | ab76442, RRID: AB_1524535 |
| Wfs1 | Proteintech | 11558-1-AP, RRID: AB_2216046 |
| Chemicals, peptides, and recombinant proteins |  |  |
| Neurobiotin tracer | Vector Laboratories | #SP-1120; RRID: AB_2313575 |
| Isoflurane inhalation anesthesia | Teva Ltd. | https://products.tevauk.com/p/Catagory?id=185 |
| Vetergesic (0.3 mg/ml Buprenorphine) | Ceva |  |
| Marcaïn (Bupivacaine) | Aspen | #PS09728 |
| Refobacin bone cement | Zimmer Biomet | #3003940002-3 |
| Tetric EvoFlow dental cement | Ivoclar Vivadent | A1 |
| DiI tracer | Life Technologies | #V22889 |
| DiO tracer | Life Technologies | #V22889 |
| R-Carvone | Sigma-Aldrich | #124931 |
| Normal horse serum | Vector Laboratories | S-2000 |
| Normal goat serum | Vector Laboratories | S-1000 |
| Cold Water Fish Skin Gelatin | Aurion | #900.033 |
| Enhancement conditioning solution ECS | Aurion | Lot 01207/3 |
| Silver enhancement solution SE-LM | Aurion | #500.022 |
| Diaminobenzidine | Sigma-Aldrich | D5637-1G |
| Critical commercial assays |  |  |
| Vectastain ABC Elite kit | Vector Laboratories | Cat# PK6100;<br>RRID: AB_2336819 |
| Vectashield Antifade Mounting Medium | Vector Laboratories | Cat# H-1000;<br>RRID: AB_2336789 |
| Experimental models: Organisms/strains |  |  |
| C57BL6j mice | Charles River Laboratories | https://www.criver.com/ |
| THY-Tau22 mice | Schindowski et al.<sup>9</sup> | http://www.informatics.jax.org/allele/MGI:3717232 |
| Software and algorithms |  |  |
| Fiji | ImageJ | https://imagej.net/software/fiji/ |
| Mathematica | Wolfram Research | https://www.wolfram.com/mathematica/ |
| MATLAB | Mathworks | www.mathworks.com/products/matlab.html |
| Zen 2008, Zen Black, Zen Blue, Axiovision | Zeiss | www.zeiss.co.uk |
| Spike2 | Cambridge Electronic Design | http://ced.co.uk/ |
| Kilosort2 | Stringer et al.<sup>84</sup> | https://github.com/MouseLand/Kilosort |
| phy | Rossant et al.<sup>85</sup> | https://klusta.readthedocs.io/en/latest/ |
| Python |  | https://www.python.org/ |
| ezTrack | Pennington et al.<sup>87</sup> | https://github.com/denisecailab/ezTrack |
| EMD | Quinn et al.<sup>40</sup> | https://gitlab.com/emd-dev/emd |
| SPSS | IBM | https://www.ibm.com/products/spss-statistics |
| Prism 7 | Prism 7 GraphPad | https://www.graphpad.com/ |
| Other |  |  |
| Buzsaki16-A16 silicon probe | NeuroNexus | https://neuronexus.com/ |
| A1x16 linear silicon probe | NeuroNexus | https://neuronexus.com/ |
| A2x2 tetrode silicon probe | NeuroNexus | https://neuronexus.com/ |

