## Supplementary Materials for "Spread of pathological human Tau from neurons to oligodendrocytes and loss of high-firing pyramidal neurons in ageing mice"

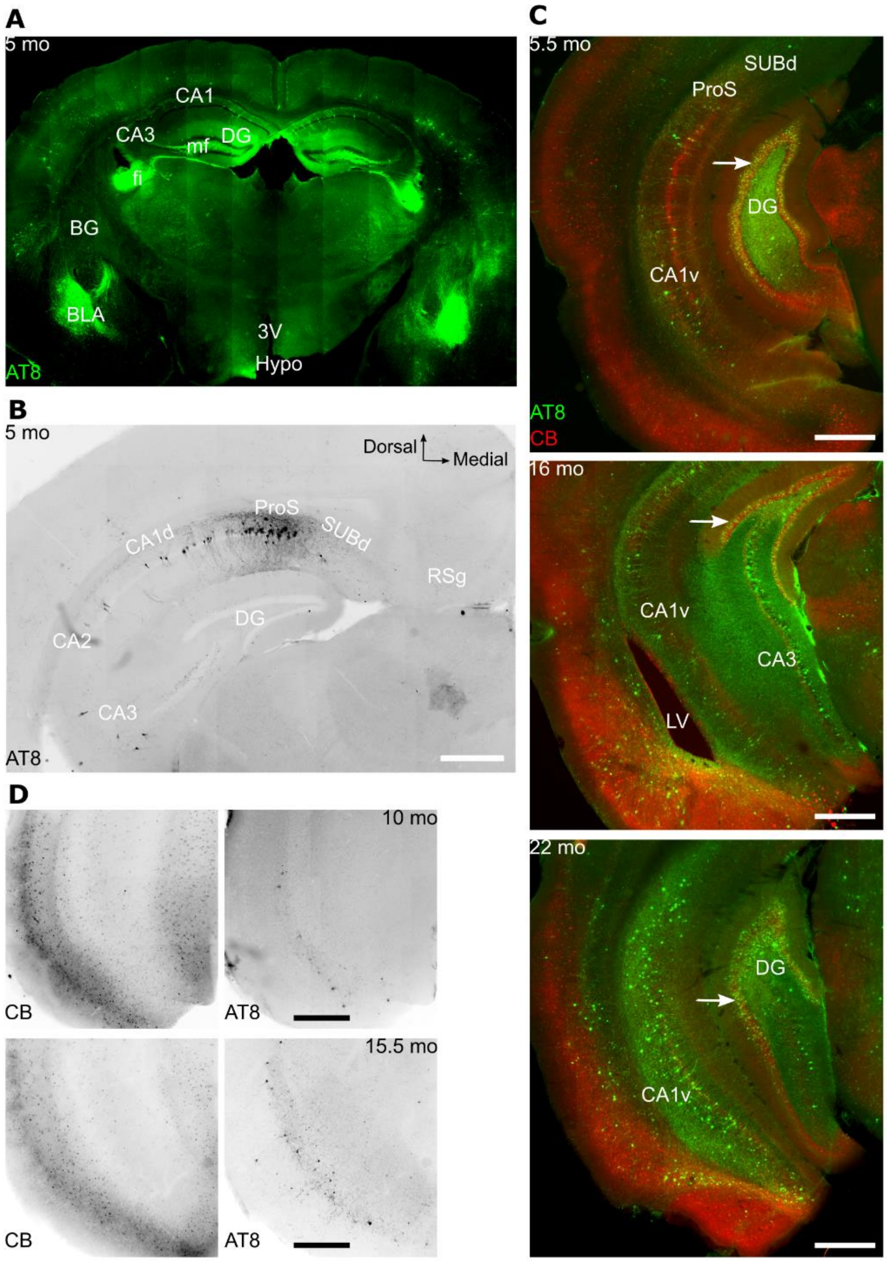


**Figure S1. Selective distribution of pTau, Related to Figure 1**

**(A)** 70 µm thick coronal section of a tg mouse (case TT71G, female, 5 mo) immunoreacted for AT8 (epifluorescence tile). Note similar distribution in both hemispheres. The lack of detectable AT8-immunoreactive granule cell somata of the dentate gyrus (DG) but immunolabelled mossy fibers (mf) suggests that the affected granule cells are from more temporal regions of the DG. **(B)** Dense pTau localized to the ProS of a different 5 mo tg mouse (case TT71F, female, littermate of case shown in (A)). Note lack of pTau in the DG, SUBd, RSg, CA2 and most of cortex. Section more caudal than that shown in A. Inverted contrast epifluorescence. Dorsal and medial directions also apply to C and D. **(C)** Caudal sections from 3 different cases (top to bottom: TV133, male, 6 mo; TT11F, male, 16 mo; TV145, male, 22 mo) immunoreacted for pTau (AT8, green) and CB (red). Note pTau+ granule cells of the DG (arrows). **(D)** Sections of entorhinal cortex (CB and AT8, inverted contrast epifluorescence) for cases TV134 (female, 10mo, top row) and TT21E (female, 16 mo, bottom row). All scale bars 500 µm. Abbreviations: BG, basal ganglia (putamen and pallidum); Hypo, hypothalamus; 3V, third ventricle; RSg, granular retrosplenial cortex; CA1v, ventral CA1; LV, lateral ventricle.

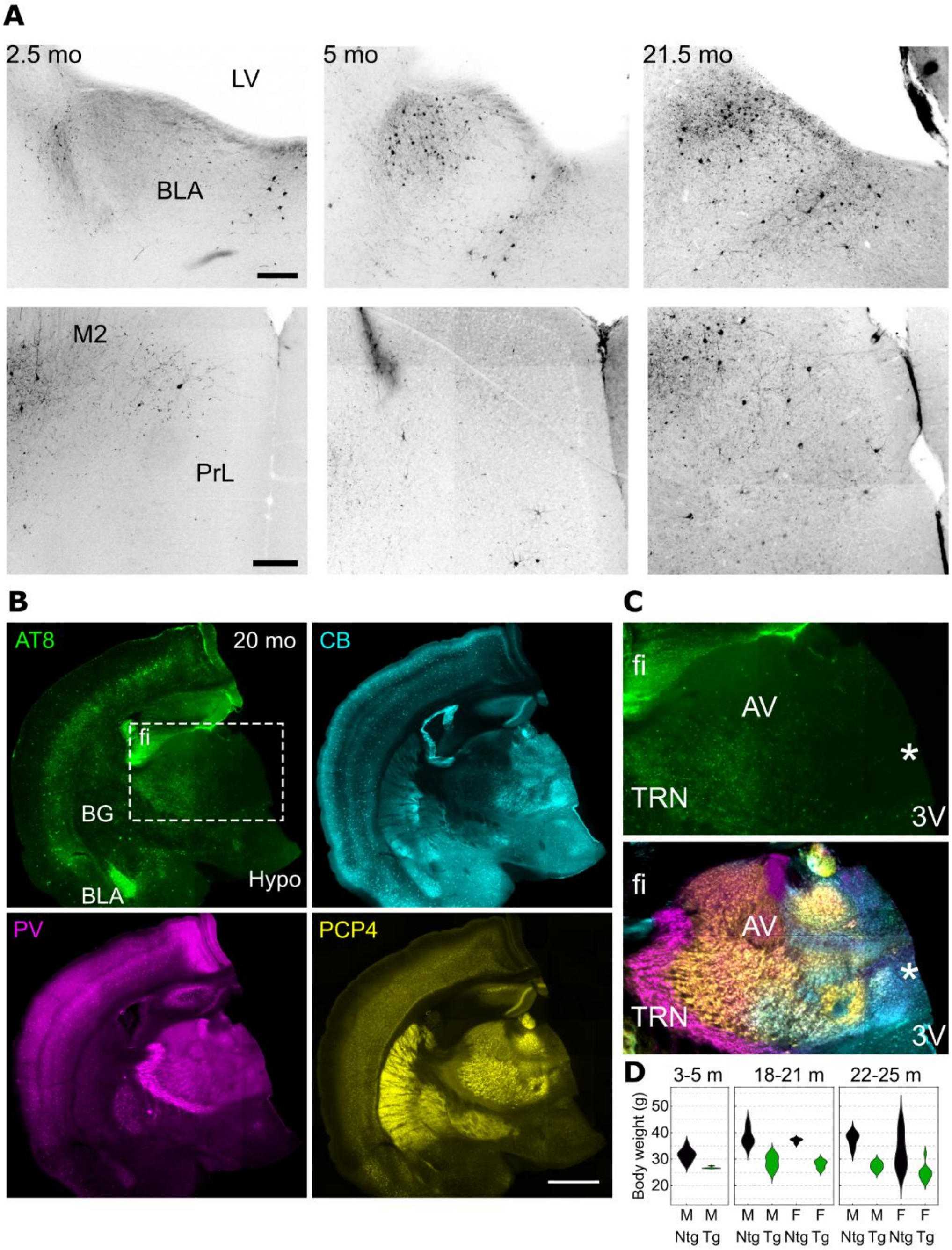


**Figure S2. Selective distribution of pTau, Related to Figure 1**

**(A)** Immunoreactivity for AT8 (pTau+) at different ages in the amygdala (top row) and medial prefrontal cortex (bottom row). Widefield epifluorescence, reverse contrast. Left, case TT114A, male, 2.5 mo; middle, case TV132, male, 5 mo; right, case TT33G, female, 22 mo. **(B)** 70 µm thick section of a 20mo tg mouse immunoreacted for AT8 (green), CB (cyan), PV (magenta) and PCP4 (yellow) (epifluorescence tile). Detail of isocortex shown in Fig. 1d. Scale bar, 1 mm. **(C)** Detail of boxed region from B, showing pTau+ axons in the fimbria (fi) and thalamic reticular nucleus (TRN) but a lack of neurofibrillary tangles and threads in the anterior thalamus (including the anteroventral nucleus, AV) and midline thalamus (asterisk). **(D)** Distribution charts of tg and ntg mouse weights at different age ranges in months (m). Abbreviations: BLA, basolateral amygdala; LV, lateral ventricle; PrL, prelimbic cortex; M2, secondary motor cortex; M, male; F, female; BG, basal ganglia (putamen and pallidum); Hypo, hypothalamus; 3V, third ventricle.

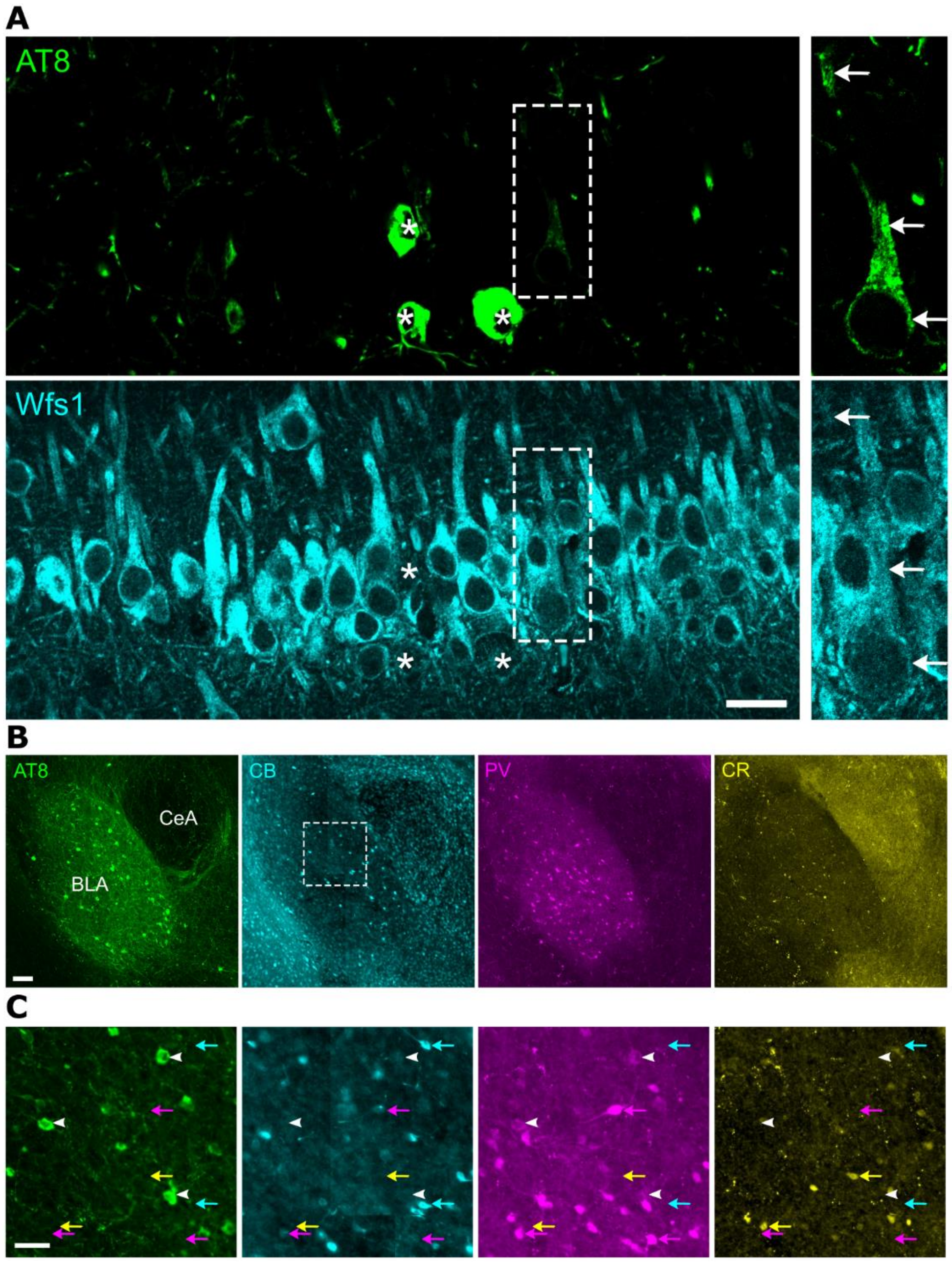


**Figure S3. Principal neurons containing pTau in the hippocampus and amygdala, Related to Figure 2**

**(A)** Wfs1 immunoreactivity (cyan) in CA1d pyramidal neurons along with few strongly pTau+ neurons (AT8, green, asterisks). Right, detail of boxed region showing a weakly pTau+ large pyramidal neuron (contrast enhanced) with weak Wfs1 immunoreactivity (arrows). Confocal single optical section (1 µm thick), case TV133 (5.5 mo). Scale bar 20 µm. **(B)** Neurons immunoreactive for pTau in the basolateral amygdala (BLA) in relation to CB (cyan), PV (magenta) and CR (calretinin). The low-level nuclear background in the cyan channel is due to autofluorescence. Note lack of pTau in central amygdala (CeA). Widefield epifluorescence tile, case TT71G (5 mo). Scale bar, 100 µm. **(C)** Detail of boxed region from B. Colored arrows highlight lack of colocalization. Note, 'rings' in the magenta channel (arrowheads) represent cross-excitation from strongly AT8 immunoreactive cells from the green channel rather than colocalization. Quantification: n=66/67 pTau+ cells in BLA lacked PV, CR and CB; n=1/67 was CB+. Scale bar, 50 µm.

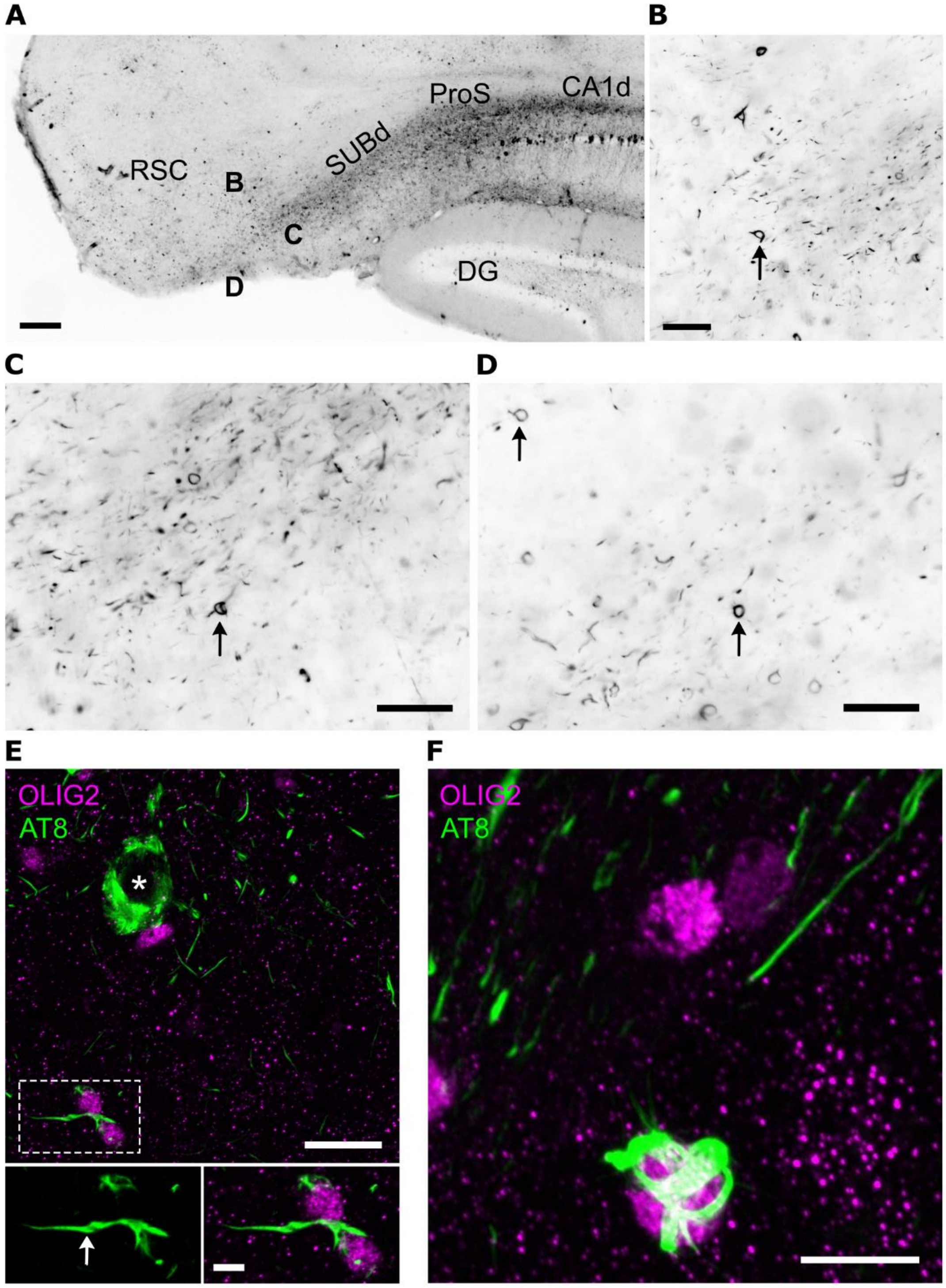


**Figure S4. Cortical pTau+ neurons and oligodendrocytes, Related to Figure 3**
**(A)** Part of the hippocampal formation (SUBd, DG, CA1d) and granular retrosplenial cortex (RSC) in an aged tg mouse (23 mo, case TT21d). Note lack of pTau+ neurons in ProS, SUBd and RSC in contrast to CA1d. Widefield epifluorescence (inverted contrast). **(B-D)** Enlargement of the areas marked in A. Arrows highlight typical coiled bodies. **(E)** Neurofibrillary tangles (asterisk, AT8, green) in a deep layer cortical neuron, which lacked immunoreactivity for Olig2 (magenta). Bottom, enlarged view of two oligodendrocytes from boxed region, indicating a long process marked by pTau (arrow). **(F)** An unusually dense coiled body surrounding an Olig2+ nucleus in the fornix. Scale bars (µm): A, 200; B-D, 50; E, 20, 5; F, 10. Confocal maximum intensity projection thickness (µm): E, 2.8; F, 4.8.

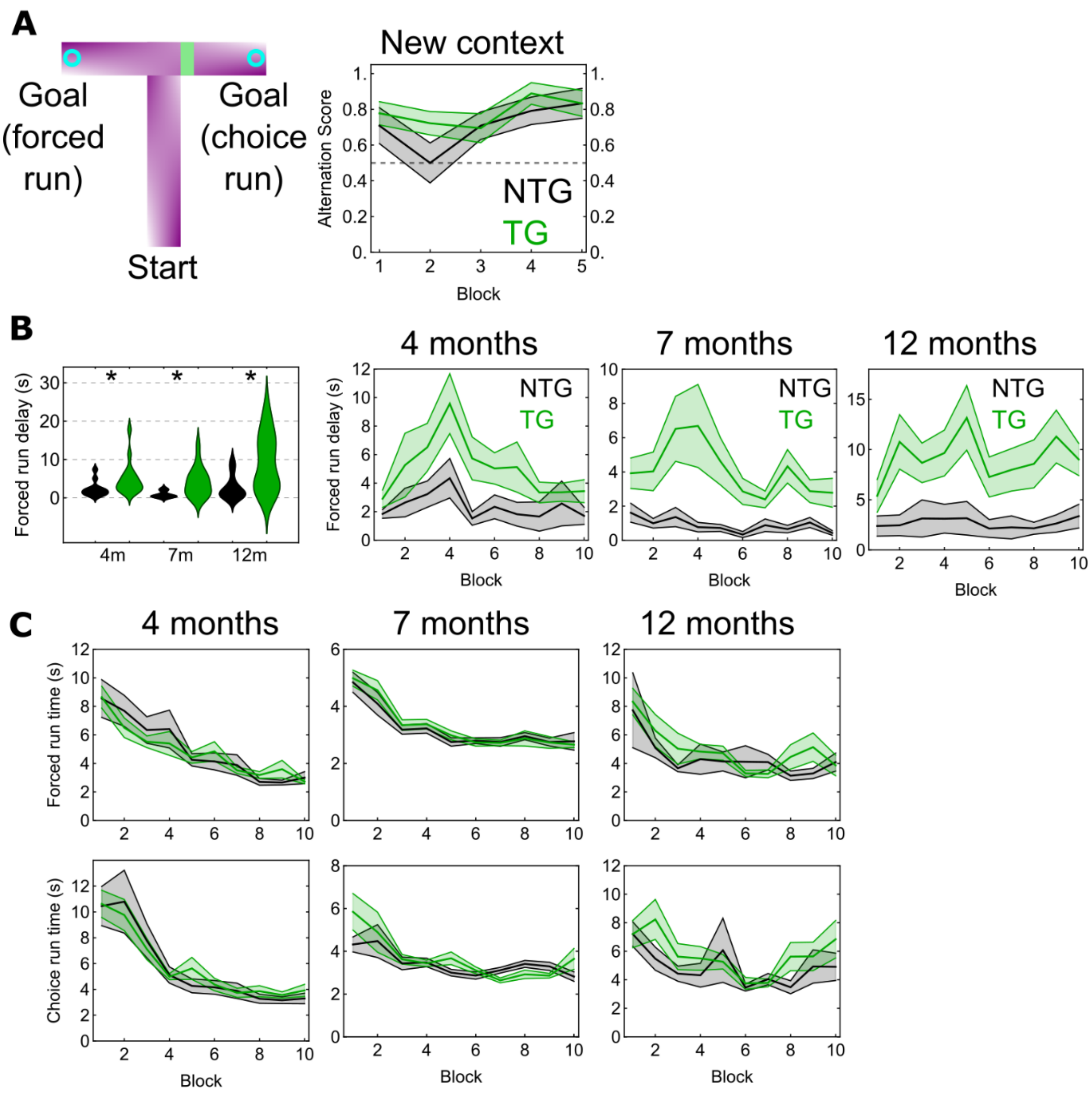


**Figure S5. Non-matching to place T-maze task, Related to Figure 4**

**(A)** Non-matching to place T-maze task but in a different, novel context for ageing (13 mo) mice (5 blocks). **(B)** Left, distribution chart of forced run delay times for the original T-maze shown in Fig. 4 (ntg, black; tg, green) during training at 4, 7, and 12 mo. Asterisks, significant difference between groups. Right, mean and standard error across blocks. **(C)** Mean and standard error of forced run times (top) and choice run times (bottom) across blocks (running from the end of the start arm to one of the goals). Note similar run times for both genotypes. Abbreviations: NTG, non-transgenic; TG, transgenic mice.

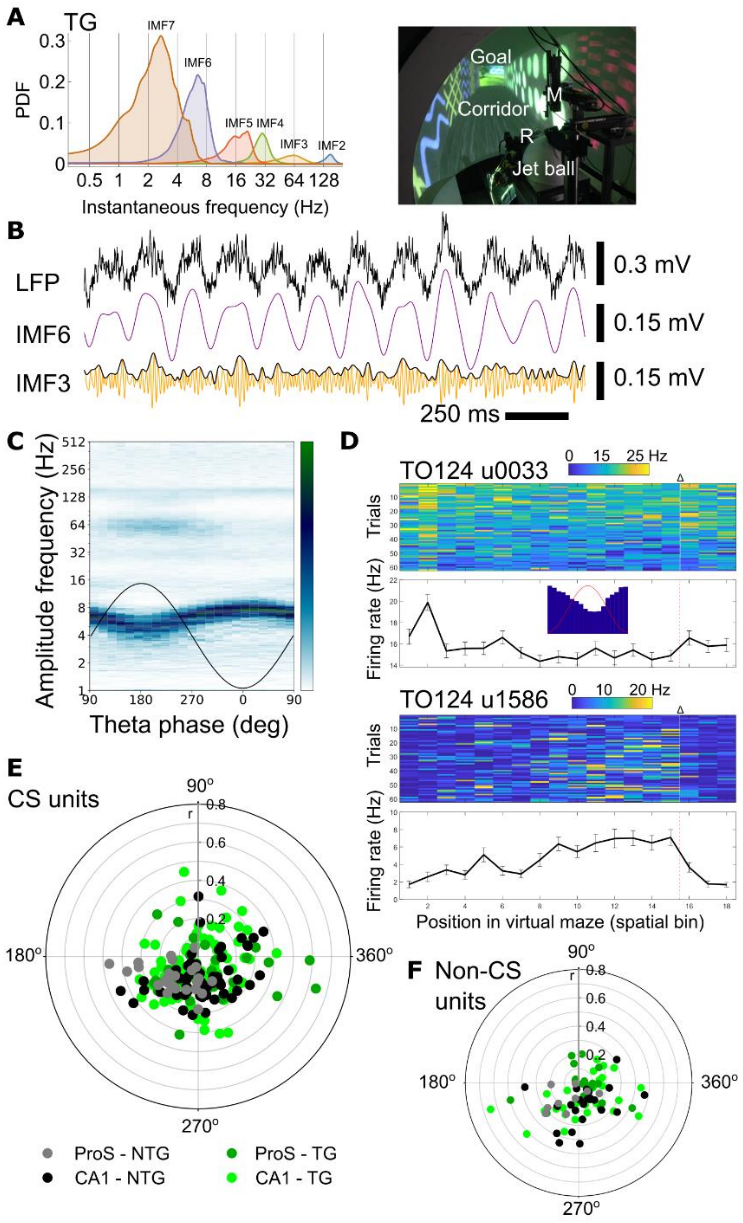


**Figure S6. Network oscillations and spike timing during goal-directed navigation, Related to Figures 5 and 7**

**(A)** Probability density function (PDF) of instantaneous frequency as in Fig. 5d, for an aged (22 mo) tg mouse (case TV145). Right, photograph of virtual environment and jet ball. A micromanipulator (M) supports the probe, and the mouse runs for a reward (R) delivered by some tubing at the goal. **(B)** Unfiltered wideband LFP from the same tg mouse as in A along with IMF6 (purple, theta) and IMF3 (orange, mid-gamma). Black curve is the instantaneous amplitude for IMF3. **(C)** Co-modulogram for the same tg mouse binned for IMF6 (theta) phase. Note phase amplitude coupling for the 'mid-gamma' range (corresponding to IMF3) on the peak to descending theta phase. Theta frequency is also modulated by phase. Black curve, sinusoidal schematic theta cycle. Color bar: green, maximum; white, minimum amplitude. **(D)** Firing rates of a non-CS unit (u0033) and a CS unit (u1586) from Fig. 7d (case TO124, tg mouse). Note drop in firing rate following the reward (Δ) for u1586. Color bars, 1 to 99 % of linearized firing rate (blue minimum, red maximum). **(E, F)** Polar plots of preferred theta phase (polar axis) with respect to coupling strength (vector length, r, radial axis) for CS units (E) and non-CS units (F). Also see Table S1.

| Site | Animal | Genotype | Age | Sessions | Trials | CS units / PCs | Non-CS units |
|---|---|---|---|---|---|---|---|
| **ProS/SUBd** | TO124 | tg | 23 | 1 | 63 | 16 | 11 |
| **ProS/SUBd** | TO125 | tg | 23 | 1 | 35 | 7 | 12 |
| **ProS/SUBd** | TV170 | tg | 19 | 4 | 411 | 35 | 15 |
| **Total** | **-** | **tg** | **-** | **6** | **509** | **58** | **38** |
| **ProS/SUBd** | TV149 | ntg | 24 | 1 | 53 | 14 | 5 |
| **ProS/SUBd** | TV153 | ntg | 20 | 1 | 44 | 21 | 5 |
| **ProS/SUBd** | TV165 | ntg | 16 | 1 | 14 | 4 | 2 |
| **Total** | **-** | **ntg** | **-** | **3** | **111** | **39** | **12** |
| **CA1** | TO124 | tg | 23 | 1 | 2 | 2 | 2 |
| **CA1** | TV145 | tg | 22 | 2 | 35 | 56 | 10 |
| **CA1** | TV179 | tg | 18 | 3 | 95 | 16 | 12 |
| **Total** | **-** | **tg** | **-** | **6** | **132** | **74** | **24** |
| **CA1** | TV149 | ntg | 24 | 1 | 82 | 11 | 3 |
| **CA1** | TO126 | ntg | 22 | 1 | 13 | 3 | 4 |
| **CA1** | TV153 | ntg | 20 | 1 | 37 | 9 | 9 |
| **CA1** | TV167 | ntg | 17 | 2 | 62 | 23 | 1 |
| **Total** | **-** | **ntg** | **-** | **5** | **194** | **46** | **17** |
| **CA1** | B148 | wt | | 1 | | 26 | - |
| **CA1** | B149 | wt | | 2 | | 16 | - |
| **CA1** | B150 | wt | | 2 | | 37 | - |
| **CA1** | B153 | wt | | 1 | | 20 | - |
| **CA1** | B154 | wt | | 1 | | 23 | - |
| **CA1** | B160 | wt | | 3 | | 39 | - |
| **Total** | **-** | **wt** | **-** | **10** | **-** | **161** | **-** |

**Table S1. Silicon probe recordings in the hippocampal formation of aged mice, Related to Figure 7**

For comparison, younger 'B' animals are from Lasztoczi and Klausberger.[1] Abbreviations: tg, transgenic; ntg, non-transgenic littermates; wt, wild type; ProS, dorsal prosubiculum; PC, putative pyramidal cells. Age is in months. Trial counts include only the trials containing reliable unit recordings, excluding periods of probe drift and the first trial of each session (n=946/1135 trials).

1. Lasztoczi, B., and Klausberger, T. (2017). Distinct gamma oscillations in the distal dendritic fields of the dentate gyrus and the CA1 area of mouse hippocampus. Brain Struct Funct *222*, 3355-3365. 10.1007/s00429-017-1421-3.
